# Reprogrammed neutrophils with impaired transit mechanics drive multi-organ capillary stalling after stroke

**DOI:** 10.64898/2026.02.10.705064

**Authors:** Jeanne Droux, Julien Husson, Chaim Glück, Hanna Preuss, Alexia Del Campo Fonseca, Talia Bergaglio, Lukas B. Otto, Colin Sparano, Ladina Hösli, Lukas Glandorf, Bruno Palmier, Sonia Tugues Solsona, Daniel Razansky, Isabelle Margaill, Melanie Greter, Daniel Ahmed, Maria Casanova Acebes, Nozomi Nishimura, Chris B. Schaffer, Andres Hidalgo, Bruno Weber, Susanne Wegener, Mohamad El Amki

## Abstract

Beyond the focal brain lesion, stroke causes systemic complications including cardiac failure, pneumonia, renal injury, and sustained immune dysfunction. The source of this multiorgan vulnerability remains unresolved. By imaging over 16,000 vessels of healthy, inflamed and ischemic brains, we identify a circulating neutrophil subpopulation reprogrammed by stroke into a pathological stalling phenotype, occluding capillaries in the brain, heart, kidneys, retina and lungs. Combining transcriptomics, genetic models, integrated microfluidics, cell mechanics assays, and *in vivo* imaging, we show that this subpopulation exhibits an atypical morphology, increased actin polymerization, and heightened adhesion that impair transit through capillary networks. This phenotype is present in patients with stroke, transmissible by adoptive transfer, and selectively sensitive to inhibition of the Src-family kinase Fgr. Both pharmacological and genetic inactivation of Fgr normalize neutrophil adhesion, reduce capillary stalls, and improve neurological recovery after stroke. These findings identify immune cell transit failure as a systemic driver of post-stroke pathology and a therapeutic target to improve both cerebral and multiorgan outcomes.

## Introduction

Stroke is the primary cause of disability worldwide ^1,2^. Despite advances in recanalization therapies, a major obstacle to recovery is the failure of microvascular reperfusion. Even after successful arterial recanalization, many downstream capillaries remain blocked, producing the “no-reflow” phenomenon ^3–5^. As a result, microcirculatory dysfunction persists, and more than half of patients still face death or major disability ^6–9^.

Beyond the focal brain lesion, stroke patients frequently develop systemic complications, including cardiac dysfunction, pneumonia, renal injury, and sustained immune alterations ^10–13^. These complications contribute to poor outcomes, yet the mechanisms linking focal brain ischemia to widespread microvascular dysfunction remain unknown.

Neutrophils, the most abundant circulating leukocytes, are among the first immune cells to respond to ischemia. Their diameter exceeds that of capillaries, forcing them to deform as they traverse the microcirculation ^14^. Any alteration in capillary diameter or changes in neutrophil deformability may transform neutrophils from flowing cells into intravascular plugs. While their inflammatory functions are well established ^15–20^, their contribution to microvascular flow disruption across organs remains poorly understood.

Several studies have demonstrated that neutrophils contribute to capillary stalls in the brain after stroke and in other inflammatory conditions, including Alzheimer’s disease, diabetes, epileptic seizures and traumatic brain injury ^21–29^. The mechanisms behind these stalls are likely multifactorial. While previous pioneering work identified that part of capillary stalling arises from local vascular damage (direct endothelial cell or pericyte dysfunction), the contribution of neutrophil programs to stalls remains unclear.

Here, we use single-cell and bulk transcriptomics, microfluidics, cell mechanics assays, and *in vivo* imaging to identify a distinct neutrophil subpopulation that emerges after stroke and exhibits a pathological stalling phenotype in the brain, heart, kidneys, retina and lungs. These cells display altered morphology, actin-driven protrusions, and increased adhesion. The phenotype is present in the patient’s blood and transmissible by adoptive transfer. Pharmacological inhibition or genetic loss of the Src-family kinase Fgr reverses the phenotype, normalizes adhesion and deformability, reduces capillary stalls, and improves neurological recovery. These findings reframe post-stroke complications as a blood-borne transport disorder and highlight neutrophil Fgr as a target to restore microcirculation dysfunction and improve multiorgan outcomes.

## Results

### Stroke induces intravascular neutrophils with a stalling behavior

Neutrophils are among the first immune cells to migrate to the injured brain, yet their dynamics within the brain vasculature remain unclear ^11,30–32^. To map intravascular neutrophil behaviors in real time, we imaged the ischemic brain vasculature by two-photon microscopy. We analyzed 7,647 neutrophils across 16,571 vessels and clustered cells based on their morphokinetic behaviors. We observed five distinct behaviors of intravascular neutrophils: flowing, crawling, adhering, squeezing, and stalling (**Figure 1A-E**, **Movies S1-S3**). Even in the same vessel, neutrophil behaviors were heterogeneous. For example, some neutrophils were flowing while others were rolling or adherent (**Figure 1A**). In capillaries, neutrophil behaviors were restricted to squeezing and stalling. Under physiological conditions, flowing and squeezing clusters represented the majority of blood circulating neutrophils. Occasionally, we observed a small cluster of stalling neutrophils (4.9%) leading to stalls in 0.37% of the total capillaries (**Figure 1C,F**). This is in line with previous studies showing that a small fraction of brain capillaries exhibits stalling even in physiological states ^23,33–35^.

**Figure 1.**
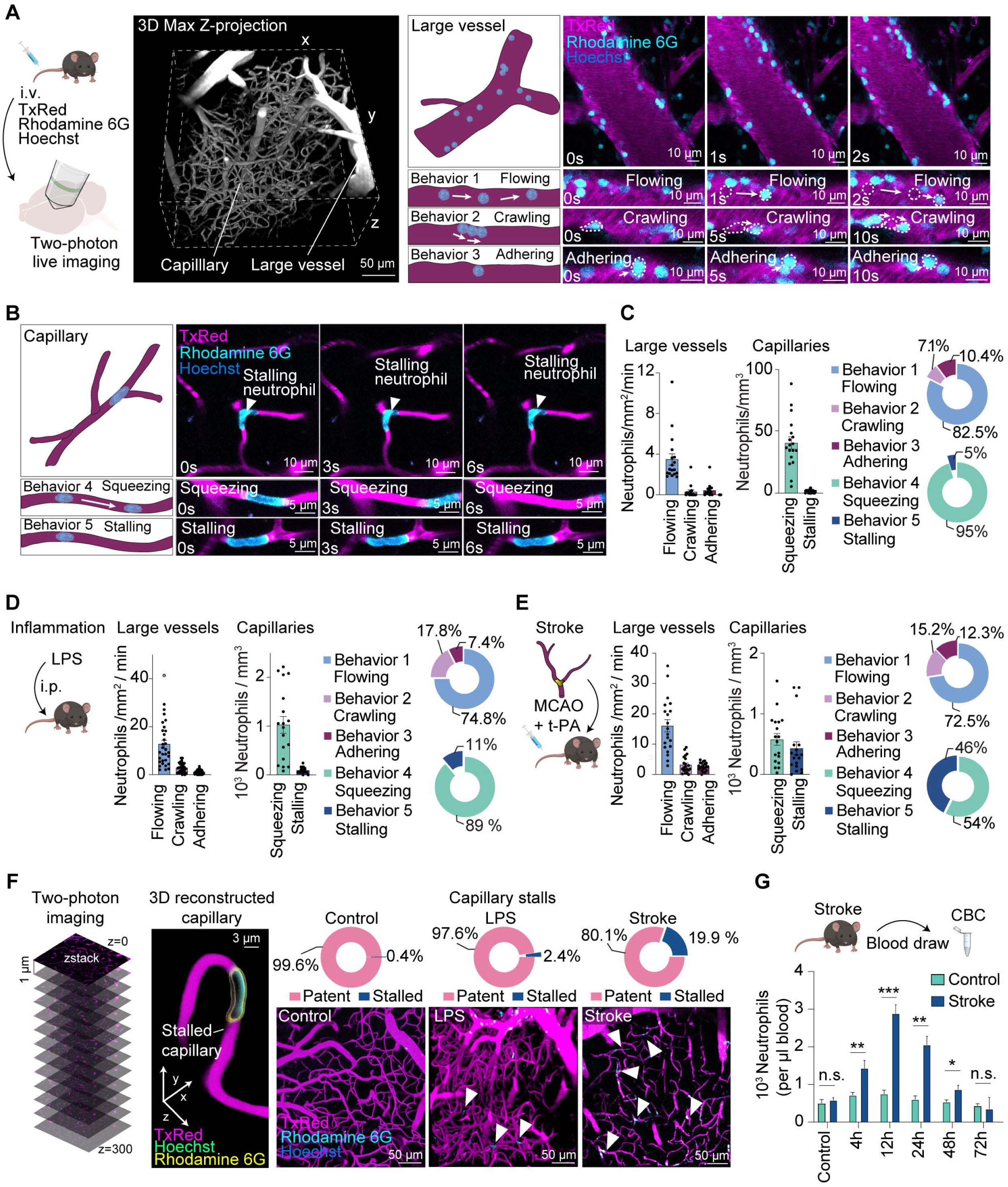
Intravascular neutrophil flow behaviors change after stroke. (**A**) Experimental design of the in vivo imaging set-up. The vascular network was labeled by intravenous (i.v.) injection of Texas Red (TxRed), neutrophils were labeled with Rhodamine 6G and cell nuclei were labeled with Hoechst. Z-projection reconstructed image of the vasculature acquired by in vivo two-photon microscopy. The representative images on the right show three different neutrophil behaviors in large vessels: flowing, crawling, and adhering. (**B**) Neutrophil behaviors in capillaries. Representative two-photon microscopy images of neutrophils in capillaries showing squeezing or stalling behavior. (**C**) Quantification of neutrophil behaviors in large vessels and capillaries of control mice (n = 37 stacks from 4 mice). On the right, percentages of the different behaviors are shown as a pie chart. (**D**) Experimental design for intraperitoneal (i.p.) LPS injection and in vivo two-photon imaging. Quantification of the flowing behaviors of neutrophils at 12 h after LPS injection (n = 40 stacks from 3 mice). (**E**) Schematic illustration of the stroke model. Mice received i.v. rt-PA infusion 30 min following stroke. Quantifications of neutrophils flow behavior at 2 h post-stroke (n = 37 stacks from 3 mice). On the right, percentages of the different behaviors are shown as a pie chart. (**F**) Three-dimensional (3D) view of a stalled capillary with a neutrophil. Representative images of capillary networks z-projected and quantifications of capillary stalls from controls (19 stacks from three mice), LPS-injected (18 stacks from three mice) and stroke mice (18 stacks from three mice). (**G**) Complete blood count (CBC) measurements of blood neutrophils in control (n = 3) and stroke (n = 3) mice over 72 h. Data are represented as scatter bar plots with the mean (± SEM), representing single stacks acquired with two-photon microscopy as black dots or bar blots with mean (± SEM). Statistical significance is reported as: n.s. P > 0.05, *P < 0.05,**P < 0.01, ***P < 0.001, ****P < 0.0001.

We next examined the changes in neutrophil behaviors in response to neuroinflammation and stroke. To mirror ischemic stroke in preclinical models, we performed thrombin injection into the middle cerebral artery (MCA) followed by thrombolysis with recombinant tissue plasminogen activator (rt-PA) at 30 min, a fibrinolytic drug that allows clot dissolution and recanalization ^36–39^. We observed a dramatic change in the neutrophil landscape, with stalling clusters increasing from 4.9% to 11% after lipopolysaccharide-induced (LPS) neuroinflammation and to 46% following stroke, triggering stalls and flow blockade in nearly half of the capillaries within the ischemic area (**Figure 1E,F**). Capillary stalls started to increase immediately after stroke and persisted until 24 hours later (**Supplemental Figure 1**). Quantifications of circulating neutrophils revealed a marked increase (up to 5-fold) starting at 4h post-stroke for up to 72h post-stroke, followed by a decline after 24h and finally normalizing by 72h after stroke onset (**Figure 1G**). These data reveal a striking, rapid, and transient behavioral shift in blood circulating neutrophils toward a flow-blocking phenotype after stroke.

### Capillary stalls extend beyond the ischemic brain

Because neutrophils heavily and preferentially infiltrate the ischemic area ^40–44^, but can also return to the circulation ^45–47^, we asked whether capillary stalling by neutrophils was restricted to the infarct. *In vivo* two-photon microscopy revealed that capillary stalls by neutrophils significantly increased in the contralateral hemisphere of stroke mice but not control mice (**Figure 2A**). Intriguingly, intracellular adhesion molecule 1 (ICAM-1) and vascular cell adhesion molecule 1 (VCAM-1), the main adhesion molecules involved in neutrophil attachment to vessel walls ^48–50^, were absent on the contralateral capillaries where capillary stalls occurred (**Figure 2B**), suggesting an active and widespread expansion of the behavioral stalling cluster leading to remote capillary stalls independently of the vascular bed.

**Figure 2.**
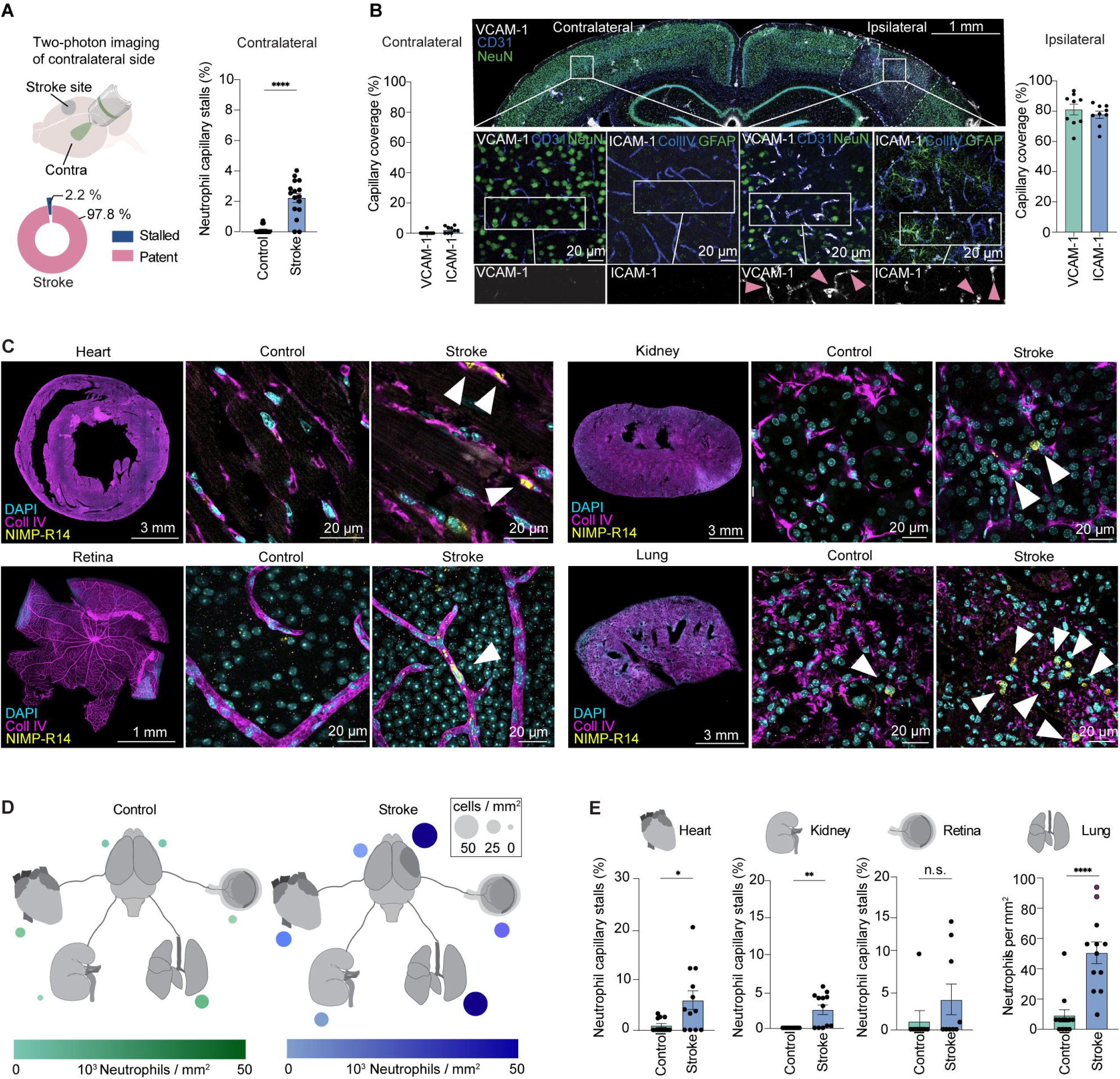
Stroke results in capillary stalls by neutrophils on the contralateral side and in peripheral organs. (**A**) Experimental design of contralateral two-photon imaging and quantification of capillary stalls by neutrophils on the contralateral side in stroke (n = 18 stacks from 4 mice) and control mice (n = 18 stacks from 4 mice, P < 0.0001, unpaired two-tailed t-test). (**B)** Immunohistochemical (IHC) staining (VCAM-1, ICAM-1, CD31, Collagen IV, GFAP and NeuN) of a coronal brain section post-stroke. Quantification of VCAM-1 (n = 9 stacks from 3 mice) and ICAM-1 coverage (n = 9 stacks from 3 mice) on capillaries in the ipsilateral and contralateral hemispheres. (**C)** Representative images showing neutrophil stalling in the heart, kidney, retina, and lungs in control and stroke mice. (**D-E)** Visualization and quantification of the number of stalls in the heart, kidney, retina and lungs in stroke (n = 12 stacks per organ from 3 mice) and control mice (n = 12 stacks per organ from 3 mice, unpaired two-tailed t-test). Data are represented as scatter bar plots with the mean, dots representing stacks acquired with confocal microscopy or two-photon microscope. Statistical significance is reported as: n.s. P > 0.05, *P < 0.05, **P < 0.01, ***P < 0.001, ****P < 0.0001.

We next asked whether stalling extended to peripheral organs beyond the brain and analyzed capillary stalls across five organs. We chose to restrict our analyses to the brain, heart, kidneys, liver, lungs, and retina because of their relatively well-understood contributions to secondary stroke complications ^51–55^. Image quantification revealed a broad distribution of neutrophil-induced capillary stalls across all these organs, in addition to the brain. This included substantial increases in capillary stalls in the heart (6%), kidneys (2.5%), lungs (5%) and retina (4%, **Figure 2C-E**). By contrast, control mice showed no stalls in these organs. Taken together, our findings suggest that stroke does not simply trap neutrophils locally but reprograms a fraction of these cells systemically into a stalling state that obstructs capillaries throughout the body.

### Stroke imprints pathogenic features in neutrophils

To corroborate the notion that stroke imprinted a stalling program in blood circulating neutrophils, we adoptively transferred neutrophils isolated from the blood of either stroke or control mice into naïve recipients and profiled their intravascular behaviors (**Figure 3A,B**).

**Figure 3.**
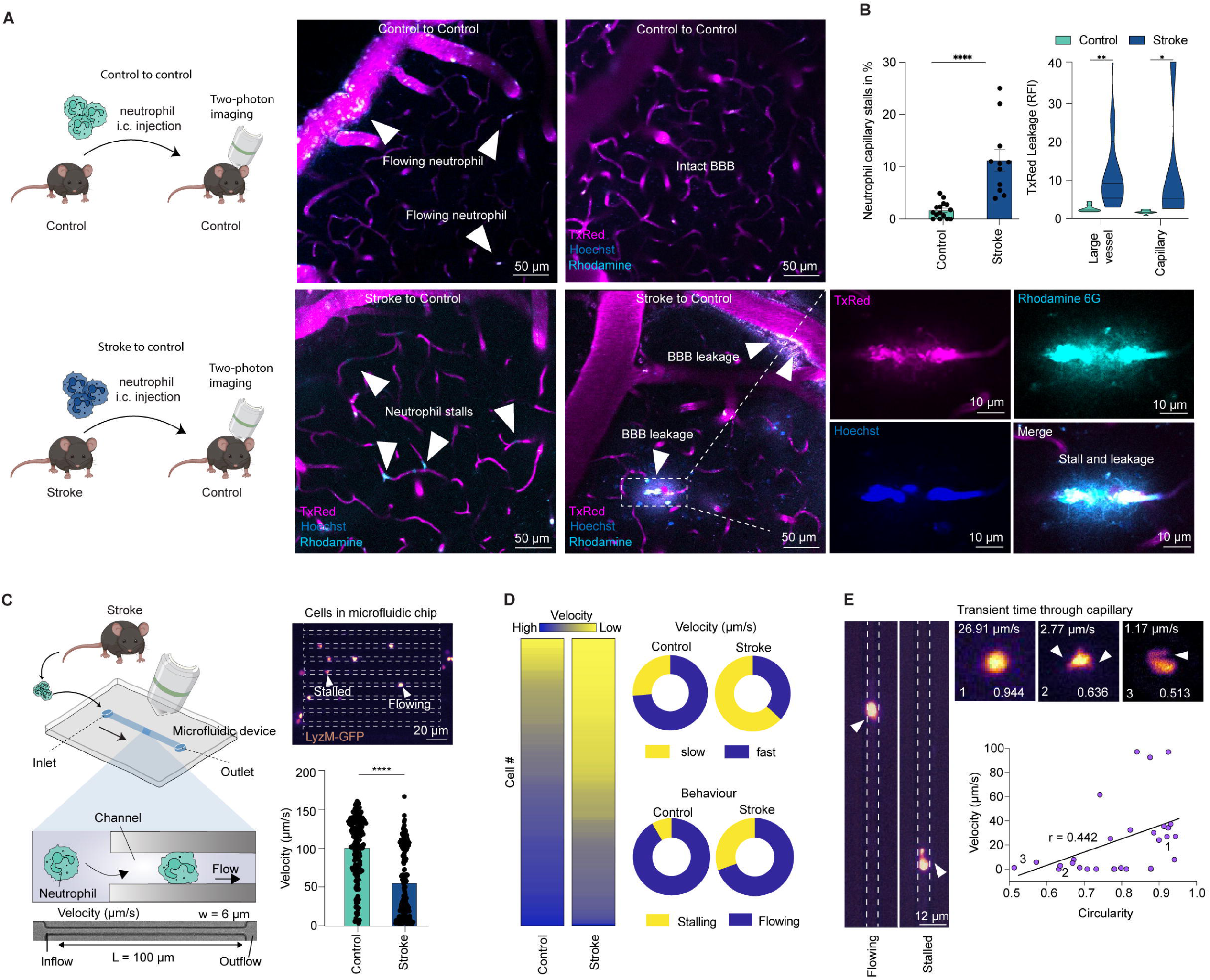
Stroke imprints a pathogenic feature in neutrophils observed after adoptive cell transfer and in microfluidic devices. (**A**) Schematic for adoptive cell transfer experiments. Representative two-photon microscopy images from control mice injected with control-derived and stroke-derived neutrophils illustrating capillary stalls and BBB leakage. (**B**) Quantification of neutrophil-induced capillary stalls in healthy receivers injected with control-derived (control to control, n = 12 stacks from 4 mice) and stroke-derived neutrophils (stroke to control, n = 11 stacks from 4 mice, P < 0.0001, unpaired two-tailed t-test). On the right, quantifications of relative TxRed fluorescence intensity in brain tissue after adoptive cell transfer. Quantifications were performed to illustrate leakage around large vessels (control to control, n = 9 stacks from 4 mice and stroke to control, n = 9 stacks from 4 mice; unpaired two-tailed test) and capillaries (control to control, n = 10 stacks from 4 mice, and stroke to control, n = 5 stacks from 4 mice; unpaired two-tailed t-test). (**C**) Scheme of microfluidic device and experimental setup to assess neutrophil flow ex vivo. Brightfield image of PDMS microfluidic chips showing neutrophils flowing through capillary channels. On the right, quantifications of neutrophil velocity from controls (n = 191 cells from 6 mice) and stroke mice (n = 212 cells from 5 mice). (**D**) Heat map of the differentially scored velocities of neutrophils with high (blue) and low velocities (yellow) illustrating the heterogeneity of neutrophil behaviors in control and stroke mice. The right upper panel represents percentages of high and slow flowing cells. The lower panel represents stalling and flowing behaviors of neutrophils in microfluidic capillary channels. (**E**) Representative images of flowing and stalling neutrophils in microfluidic capillaries. Right upper images show single neutrophils with different velocities in the capillary and their corresponding circularity coefficient assessed before entering microfluidic channel. On the right, correlation of neutrophils velocities and their morphology (r = 0.441, P = 0.0164). Data are represented as scatter bar plots with the mean (± SEM), as violin plots showing the median and upper and lower quartiles, and scatter plots with correlation line. Statistical significance is reported as: n.s. P > 0.05, *P < 0.05, **P < 0.01, ***P < 0.001, ****P < 0.0001.

To specifically isolate neutrophils and confirm their identity, we used flow cytometry or gradient centrifugation protocols achieving 95-98% purities ^56,57^. 100,000 neutrophils from control and stroke-induced mice were injected into the internal carotid artery to force passage through the brain capillary network, and live imaging with two-photon microscopy revealed that transferred neutrophils from both groups were detected in the brain vascular network of recipient mice, indicating successful transfer (**Figure 3A,B**). Notably, while control neutrophils did not trigger capillary stalls (**Figure 3A**), stroke-derived neutrophils induced stalls in 10% of the capillaries of naïve mice (**Figure 3B**). When the total number of injected neutrophils was increased to 300,000 cells, we observed that stroke-derived neutrophils triggered blood–brain barrier (BBB) leakage. This was characterized by extravasation of fluorescent plasma dye, nuclear stain, and neutrophil migration into the affected area (**Figure 3B**). Together, these data show that stroke imprints a stalling and pathogenic program in circulating neutrophils that is sufficient to block healthy capillaries. These findings also suggests that stroke imprints a distinct pathogenic program in neutrophils even in the absence of overt vascular injury in the recipient.

### Morphological landscape of stalling neutrophils

Next, we investigated the cellular basis of this pathogenic program. First, we developed a microfluidic system in which neutrophils were perfused into channels mimicking the narrow capillaries of the brain. In this setup, cells were introduced with a syringe connected to a pump into non-adhesive polydimethylsiloxane (PDMS) channels (6 µm diameter). Cells were introduced at an initial constant velocity, allowing for the study of cell-intrinsic changes and flow behaviors. Using high-speed imaging, we tracked neutrophils isolated from stroke-induced and control *LyzM*^GFP^ reporter mice (**Figure 3C**). Automated tracking of individual cells revealed a bimodal distribution of cell speed as they travelled through the channels (**Figure 3C** and **Movie S4**), suggesting the presence of two behaviorally different subpopulations of neutrophils (based on flow speeds below versus above 75 µm/s), mirroring the *in vivo* heterogeneity in cellular dynamics (**Figure 1**). We observed that the cluster of neutrophils with slower velocity and stalling behavior increased from 26% to 63% after stroke (**Figure 3D**).

Cellular motility has been shown to reflect morphological features ^58–60^. Therefore, we investigated whether stalling neutrophils show altered shapes, even before confinement, since passage through capillaries can deform cells. Indeed, we found that slower cells exhibited a basal altered circularity and were more likely to stall (**Figure 3E**). We refined these analyses of neutrophil shapes using live-cell imaging and immunofluorescence of the cytoskeleton. Morphological analysis revealed an atypical neutrophil subpopulation with reduced circularity and increased size (**Figure 4A-C**). Isolated neutrophils were then stained for DNA and intracellular F-actin, a key regulator of cellular motility ^61–63^. Confocal microscopy revealed that atypical neutrophils displayed a thick actin-polymerized cortex and a cell membrane rich in protrusions. In contrast, neutrophils from controls were mostly small and round, which we refer to as “conventional cells” (**Figure 4D**). We then mapped F-actin dynamics in neutrophils *in vivo* by imaging *LifeAct*^eGFP^ reporter mice by two-photon microscopy. In large vessels, we observed distinct actin dynamics profiles in neutrophils depending on their flow behavior; flowing cells showed a thin cortex and reduced actin intensity, while adhering cells showed high actin polymerization (**Supplemental Figure 5, Movie S5**). In capillaries, we observed that flowing neutrophils exhibited F-actin flow with a high back-to-front F-actin ratio (**Figure 4F,G**). In contrast, stalling neutrophils exhibited a significant decrease in the back-to-front F-actin ratio and increased overall F-actin polymerization, resulting in low actin flow (**Figure 4G, Movie S6**). Images of *LifeAct*^eGFP^ expression at sequential time points show the marked differences in F-actin distribution in flowing versus stalling cells (**Figure 4F,G**). Together, our data indicate that the pathogenic program in circulating neutrophils features altered morphology and F-actin dynamics in atypical neutrophils.

**Figure 4.**
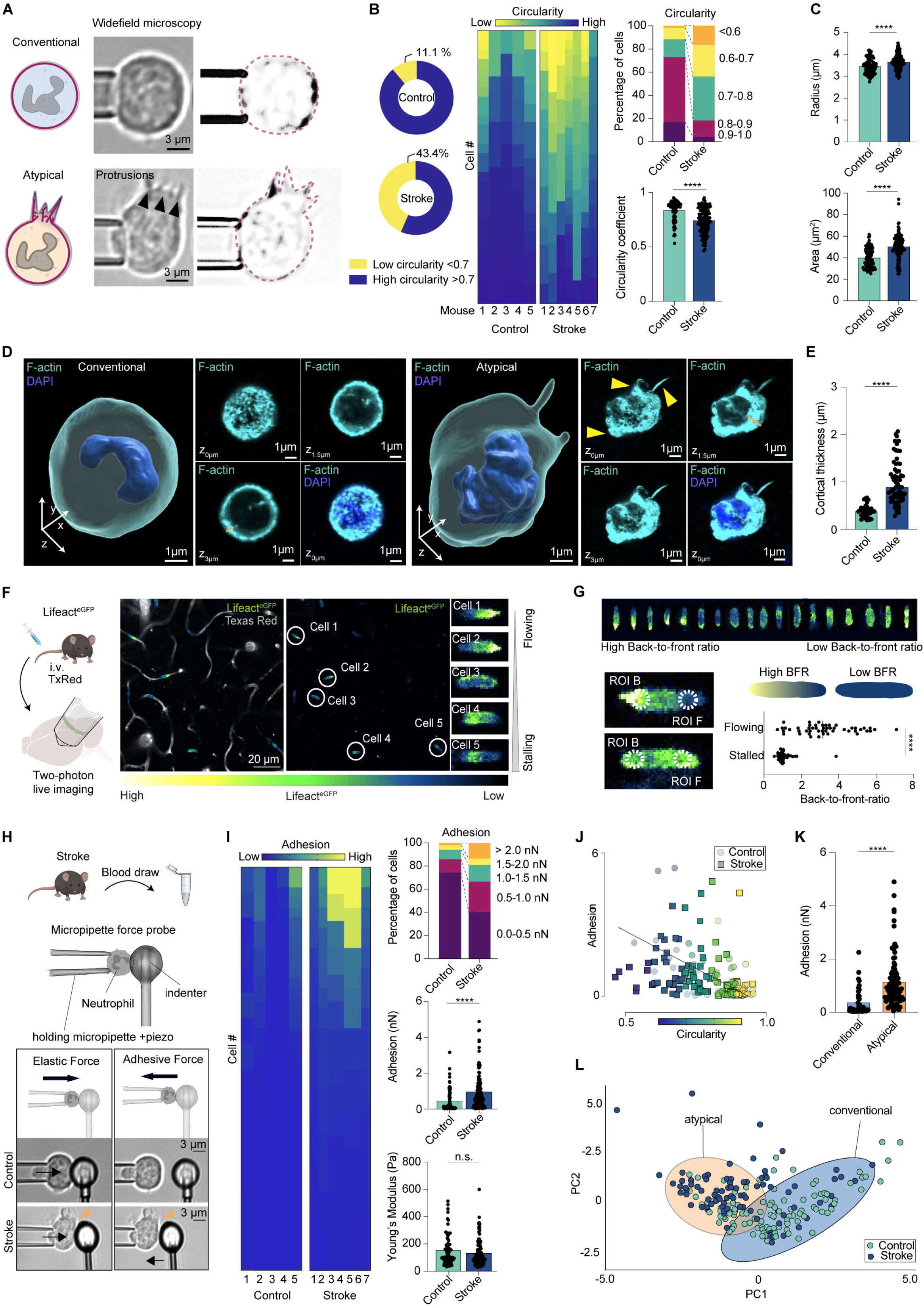
Stroke reprograms an atypical neutrophil subpopulation with morphological and cell mechanical alterations. (**A**) Schematic and representative images showing the shape of conventional and atypical neutrophils acquired with widefield microscopy. (**B**) Left, pie chart displaying the proportion of neutrophils with high and low circularity (threshold 0.7) in control and stroke mice. Heat map of single cells and their cellular circularity from high (blue) to low (yellow). The right panel shows percentages of neutrophils with different circularity indices in control and stroke mice (n = 211 cells from 5-7 mice per group). Circularity was measured as 41[(area/perimeter^2^). (**C**) Quantification of neutrophil area and radius in control and stroke mice (n = 333 cells from 5-7 mice per group, two-tailed Mann-Whitney test). (**D**) Representative images of immunohistochemical staining and 3D reconstruction of conventional and atypical neutrophils. stained with F-actin and DAPI. (**E**) Quantification of actin thickness in neutrophils (n = 108 cells from three mice per group, two-tailed Mann-Whitney test). (**F**) Schematic of in vivo two-photon imaging of actin dynamics in neutrophils. Representative two-photon images showing LifeAct^eGFP^ (actin, green) neutrophils from Lifeact^eGFP^ mice illustrating actin polymerization in flowing and stalling neutrophils. (**G**) Quantification of the actin back-to-front ratio in stalling and flowing neutrophils (n = 91 cells in 4 mice per group, unpaired-two-tailed t-test). (**H**) Scheme of micropipette force probe measurements to assess elastic and adhesion forces in neutrophils. The lower panel shows representative images of control and stroke-derived neutrophils during force probe measurement. (**I**) Heat map of single cells and their adhesion from high (yellow) to low (blue). Right upper panel, percentages of neutrophils with different adhesion forces. The panels below show adhesion and elasticity forces of neutrophils from control and stroke mice (n = 211 cells from 5-7 mice per group). (**J**) Correlation between neutrophil adhesion and circularity (r = -0.217, P = 0.0346). (**K**) Adhesion force according to circularity index (cut-off = 0.8), separating conventional from atypical neutrophils. Conventional cells show lower adhesion compared to atypical cells (n = 133 cells from 12 mice). (**L**) K-means clustering of neutrophil behaviors and morpho-mechanical parameters including area, radius, circularity, adhesion and elasticity. Data are represented as scatter bar plots, or scatter plots representing single cells as dots. Statistical significance is reported as: n.s. P > 0.05, *P < 0.05, **P < 0.01, ***P < 0.001, ****P < 0.0001.

### Altered mechanics and increased adhesion of stroke-educated neutrophils

To define the cellular basis of stalling, we tested whether actin-driven protrusions correlated with altered cellular mechanics. To address this, we directly measured cell stiffness and adhesion using micropipette force probe microscopy, as previously described ^64–66^. This method simultaneously measures cellular morphological changes and captures cellular mechanical properties ^64^ (**Figure 4H, Movie S7**), and we adapted the technique to additionally measure adhesion forces. While overall stiffness was similar, adhesion forces nearly doubled in stroke neutrophils versus controls (**Figure 4I**). Strikingly, atypical neutrophils, which contained abundant protrusions, were more adherent (**Figure 4J,K**). Clustering analysis integrating morphological and mechanical parameters segregated a distinct population with altered morphology and function, consistent with a shift toward a stalling phenotype after stroke (**Figure 4L**). Thus, this suggests that stroke imprints a mechanical phenotype that locks neutrophils into a stall-prone state.

### Transcriptomic profiling identifies Fgr signaling during neutrophil reprogramming

We next examined whether stroke rewired the transcriptional program of neutrophils. First, to extend our findings to humans, we analyzed publicly available single-cell RNA-seq (scRNA-seq) datasets of peripheral blood from stroke patients ^67^ (**Figure 5A-C**). After performing unsupervised clustering, we identified different neutrophil clusters based on the most variable genes, revealing divergent transcriptomic signatures. We identified a neutrophil cluster whose signature was associated with cytoskeletal remodeling and abnormal morphology (**Figure 5B,C**). Notably, the post-stroke signature aligned with a pro-inflammatory phenotype, with upregulated pathways related to cytoskeleton, trafficking, and signaling, including tubulin binding and GTPase activity. These data support stroke-induced reprogramming of neutrophils toward a stalling phenotype in both mice and humans.

**Figure 5.**
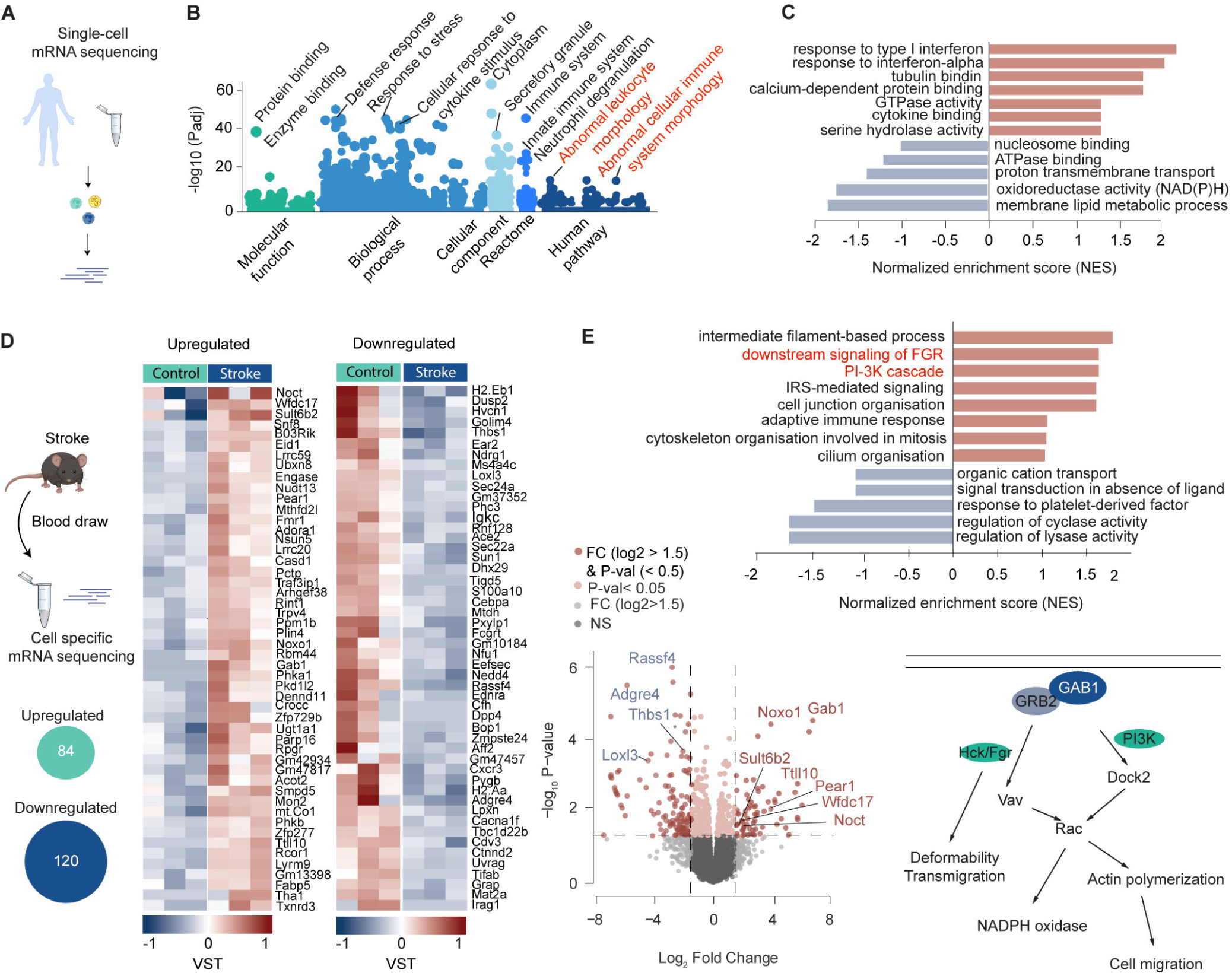
Single-cell and bulk mRNA-seq of blood neutrophils from stroke patients and mice. (**A**) Schematic of human single-cell sequencing workflow. (**B**) Gene ontology (GO) molecular function analysis of enriched genes in neutrophils of stroke patients compared to corresponding controls. GO highlights significant changes in genes responsible for molecular function, biological process, cellular component, Reactome and human pathway (Dataset from Xie et al. 2020). (**C**) Bar plots showing the normalized enrichment score (NES) of selected GO categories enriched within genes upregulated (red) or downregulated (blue) in neutrophils from stroke versus control patients. Pathways with increased or decreased NES are shown. (**D**) Schematic of neutrophil isolation for bulk mRNA sequencing. Heat map showing differential transcriptional profiles of control and stroke-derived blood neutrophils. Top 50 upregulated and 50 downregulated genes in stroke mice compared to control mice (DESeq2). (**E**) Bar plots showing NES of selected GO categories enriched within genes upregulated (red) or downregulated (blue) in neutrophils from stroke versus control mice illustrating increased signaling pathway controlled by Fgr and PI3K pathways. Below, volcano plot showing DEGs in neutrophils from stroke and control mice. The legend highlights upregulated (red) or downregulated (blue) transcripts, as well as genes not passing cutoff criteria (-log_10_ (Pvalue) < 0.05 and log2 fold change > 1.5). On the right, schematic of the PI3K and Fgr signaling pathways.

To define the molecular drivers of this phenotype, we performed bulk RNA-seq on sorted circulating neutrophils from stroke and control mice. Principal-component analysis (PCA) of average gene expression revealed clear separation between stroke and control samples, highlighting distinct transcriptional profiles. This signature included increased expression of *Noct* (Nocturnin), a circadian- regulated deadenylase linked to metabolic stress adaptation, and *Wfdc17* (WAP four-disulfide core domain 17), an interferon-inducible secreted protein associated with myeloid activation. Additional upregulated genes included *Gab1* (GRB2 associated binding protein 1), which amplifies growth factor signaling and regulates cytoskeletal dynamics and cell migration, *Pear1* (platelet endothelial aggregation receptor 1), a receptor enabling PI3K signal transduction, platelet aggregation and activation, and *Sult6b2* (sulfotransferase family 6B), a sulfotransferase implicated in metabolic regulation (**Figure 5D,E**). Adhesion-related transcripts were reduced, including *Thbs1* (thrombospondin 1), an extracellular matrix glycoprotein that mediates cell-to-cell and cell-to-matrix adhesion, and *Loxl3* (lysil oxidase like 3), an enzyme critical for maintaining the extracellular matrix (**Figure 5D,E**). Gene Ontology analysis highlighted pathways in motility, migration, intermediate filament organization, Fgr downstream signaling, PI3K-Akt signaling, and cytoskeletal remodeling (**Figure 5E**).

### Fgr inhibition restores neutrophil morphology and function

Among these pathways, Fgr-associated signaling was of particular interest, as Fgr is an Src-family tyrosine kinase that integrates cytoskeletal remodeling and actin polymerization ^68,69^. These processes drive the morphological alterations and increased adhesion ^70^, key features of the stalling phenotype. To test whether the observed neutrophil dysfunction in our models is Fgr-mediated, we pharmacologically administration the synthetic Fgr-inhibitor TL02-59 in mice subjected to stroke ^71,72^ (**Figure 6A**). Stroke mice were treated with rt-PA plus vehicle or rt-PA plus TL02-59 (5 mg/kg, i.v.). At 24 h post-stroke, we isolated neutrophils for mechanical and morphological assessment. To test the effect of Fgr inhibition by exclusively acting on neutrophils, we also treated the cells *ex vivo*. We found that TL02-59 reduced cell size, restored circularity, and decreased adhesion forces *in vivo* and *ex vivo* (**Figure 6B-D**). Treated cells passed through microfluidic channels faster, approaching control dynamics (**Figure 6E**). Thus, interfering with Fgr activity resolved the morphological alterations, reduced stickiness, and restored the flowing phenotype of neutrophils (**Figure 6F**).

**Figure 6.**
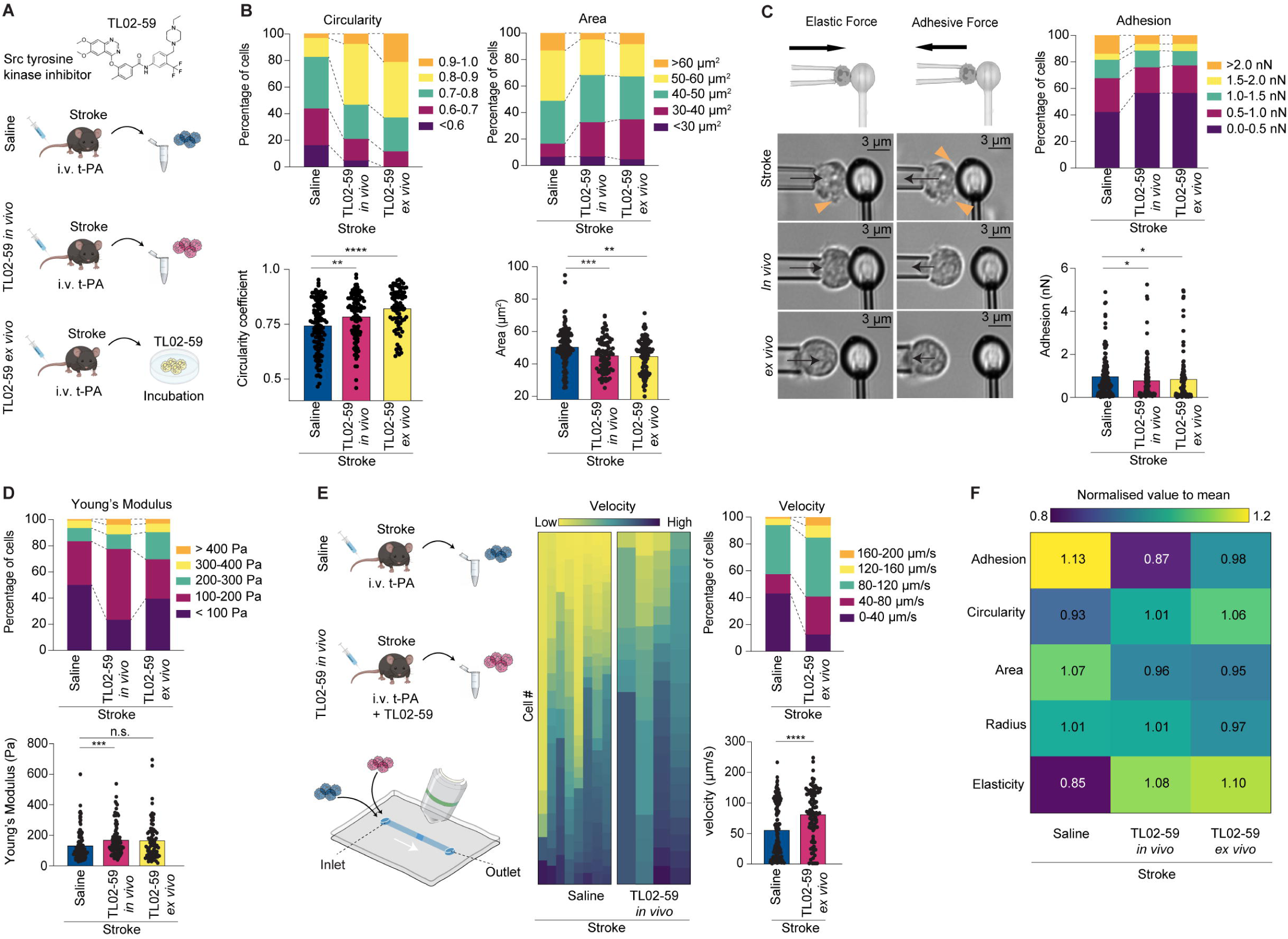
Fgr inhibition reverses morphological and behavioral alterations of neutrophils. (**A**) Experimental design for pharmacological Fgr inhibition via TL02-59 administration in vivo or ex vivo post-stroke. (**B**) Percentages of neutrophils with different circularities and areas, from stroke mice treated with saline, TL02-59 in vivo and TL02-59 ex vivo. Lower panels show circularity coefficient and area of neutrophils from stroke control compared with TL02-59 treated mice (n = 325 cells from 5-7 mice per group, Mann-Whitney test, two-tailed). (**C**) Representative images of micropipette force probe measurements of neutrophils under different treatments. On the right, percentages of neutrophils with various adhesion forces. Below, adhesion force of neutrophils from stroke control, stroke TL02-59 ex vivo-treated and TL02-59 in vivo-treated mice (n = 325 cells from 5-7 mice per group, Mann-Whitney test, two-tailed t). (**D**) Percentages of neutrophils with different Young’s modulus and Young’s modulus changes in the different stroke groups (n = 325 cells from 5-7 mice per group, Mann-Whitney test, two-tailed). (**E**) Experimental design to assess neutrophil flow in microfluidic chips. Heat map of single cells and their velocity in capillary channels from low (yellow) to high (blue). Right upper panel, percentages of neutrophils with different velocities. The panels below show velocity of neutrophils from stroke control and stroke TL02-59 treated mice (n = 308 cells from 5 mice per group). (**F**) Heat maps of relative changes in the differentially scored parameters for neutrophils, showing average mean values for adhesion, circularity, area, radius and elasticity per group, normalized to the overall mean (stroke, TL02-59 treatment in vivo and TL02-59 treatment in vitro). Data are represented as scatter bar plots. Statistical significance is reported as: n.s. P > 0.05, *P < 0.05, **P < 0.01, ***P < 0.001, ****P < 0.0001.

### Targeting Fgr reduces capillary stalls and improves stroke outcome

Having established that TL02-59 interferes with the stalling program, we tested whether targeting this pathway could improve stroke outcome *in vivo*. We treated stroke mice with rt-PA alone or rt-PA combined with TL02-59 starting at 30 minutes poststroke (**Figure 7A**). Laser speckle imaging conducted before and shortly after stroke induction confirmed that the occlusion-induced hypoperfusion was similar in both groups (**Supplemental Figure 6**), ensuring equivalent initial ischemia induction for a valid comparison between groups. Notably, *in vivo* two-photon imaging revealed that TL02-59 treatment dramatically reduced stalling in brain capillaries (**Figure 7A**).

**Figure 7.**
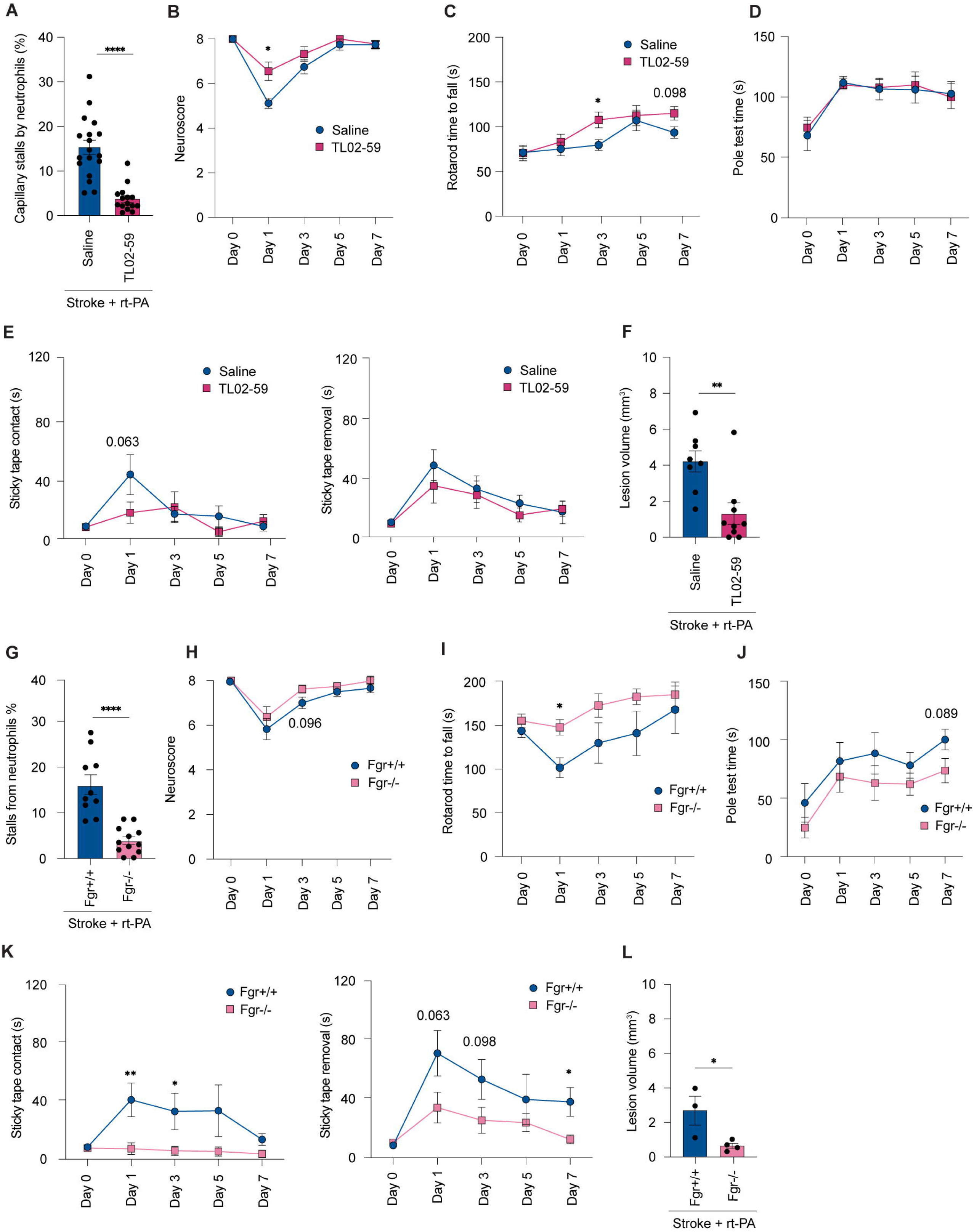
Pharmacological and genetic depletion of Fgr reduce capillary stalls and improve stroke outcome. (**A**) Quantification of capillary stalls after stroke and recanalization in control and TL02-59-treated mice (n = 34 stacks from 8-9 mice per group, P < 0.0001, unpaired, two-tailed t-test). (**B-E**) Neurological score (Neuroscore), rotarod time, pole test time and sticky tape removal assessments in saline + rt-PA-treated mice and TL02-59 + rt-PA-treated mice (n = 8-9 mice per group) at days 1, 3, and 7 after stroke.(**F**) Lesion volumes (mm^2^) at day 7 in saline + rt-PA-treated, and TL02-59 + rt-PA-treated mice (n = 8-9 per group, P<0.001, two-tailed unpaired t-test). (**G)** Quantification of capillary stalls in Fgr+/+ and Fgr-/- mice treated with rt-PA (n = 4-6 per group, P < 0.0001, unpaired, two-tailed t-test). (**H-K)** Neuroscore, rotarod time, pole test time and sticky tape removal assessment in rt-PA-treated Fgr+/+ mice and rt-PA-treated Fgr-/- mice (n = 6-8 mice per group, unpaired two-tailed t-test) at days 1, 3, and 7 after stroke. (**L)** Lesion volumes (mm^2^) at day 7 in Fgr+/+ and Fgr-/- mice treated with rt-PA (n = 3-4 mice per group, unpaired two-tailed t-test). Statistical significance is reported as: n.s. P > 0.05, *P < 0.05, **P < 0.01, ***P < 0.001, ****P < 0.0001.

To evaluate whether these TL02-59-evoked flow improvements ameliorated infarct size following stroke, we used the chemical stain 2,3,5-triphenyltetrazolium chloride (TTC), which stains live but not infarcted tissue. TTC staining 7 days after the induction of cerebral ischemia showed that TL02-59-treated mice exhibited marked reductions in infarct volume. To delineate whether reduced infarct volume translated into better neurological function, we employed several behavior tests at baseline and days 1, 3, and 7 (**Figure 7B-E**). Neurological recovery was improved across multiple behavioral assays in the TL02-59-treated mice, indicating that the cerebral function in the rescued tissue was preserved. For example, in the rotarod test, time to fall was significantly higher in TL02-59-treated mice. In the sticky tape test, control treated mice tended to show longer latencies to sense and remove the tape compared to TL02-59 treated mice, reflecting a worse neurological outcome (**Figure 7D**).

We confirmed these results using a genetic model. We generated mice lacking Fgr in leukocytes by bone marrow transplantation (**Supplemental Figure 4**), and after engraftment we subjected them to cerebral ischemia. Consistent with our TL02-59 results, Fgr-/- mice showed significantly reduced infarct volumes and neurological deficits compared to control transplanted mice (**Figure 7F-J**).

Collectively, these findings identify Fgr as a molecular driver of the pathogenic stalling behavior induced by stroke, and demonstrate the potential value of inhibiting this behavioral transition to improve systemic and neurological stroke outcomes.

### Atypical neutrophils are present in stroke patients

To examine whether our preclinical findings also translate to humans, we tested if blood from stroke patients also contains a population of neutrophils with atypical shapes. To address this, we isolated neutrophils from the blood of three healthy human donors and four stroke patients for morphological analysis (**Figure 8A**). To minimize possible confounding factors from *ex vivo* manipulation, we employed label-free digital holo-tomographic microscopy (DHTM), uses low-power laser interference to generate three-dimensional refractive index (RI) maps at nanoscale resolution, enabling quantitative morphometric analysis of live cells in aqueous environments without drying or fluorescent labeling (**Figure 8B,D**).

**Figure 8.**
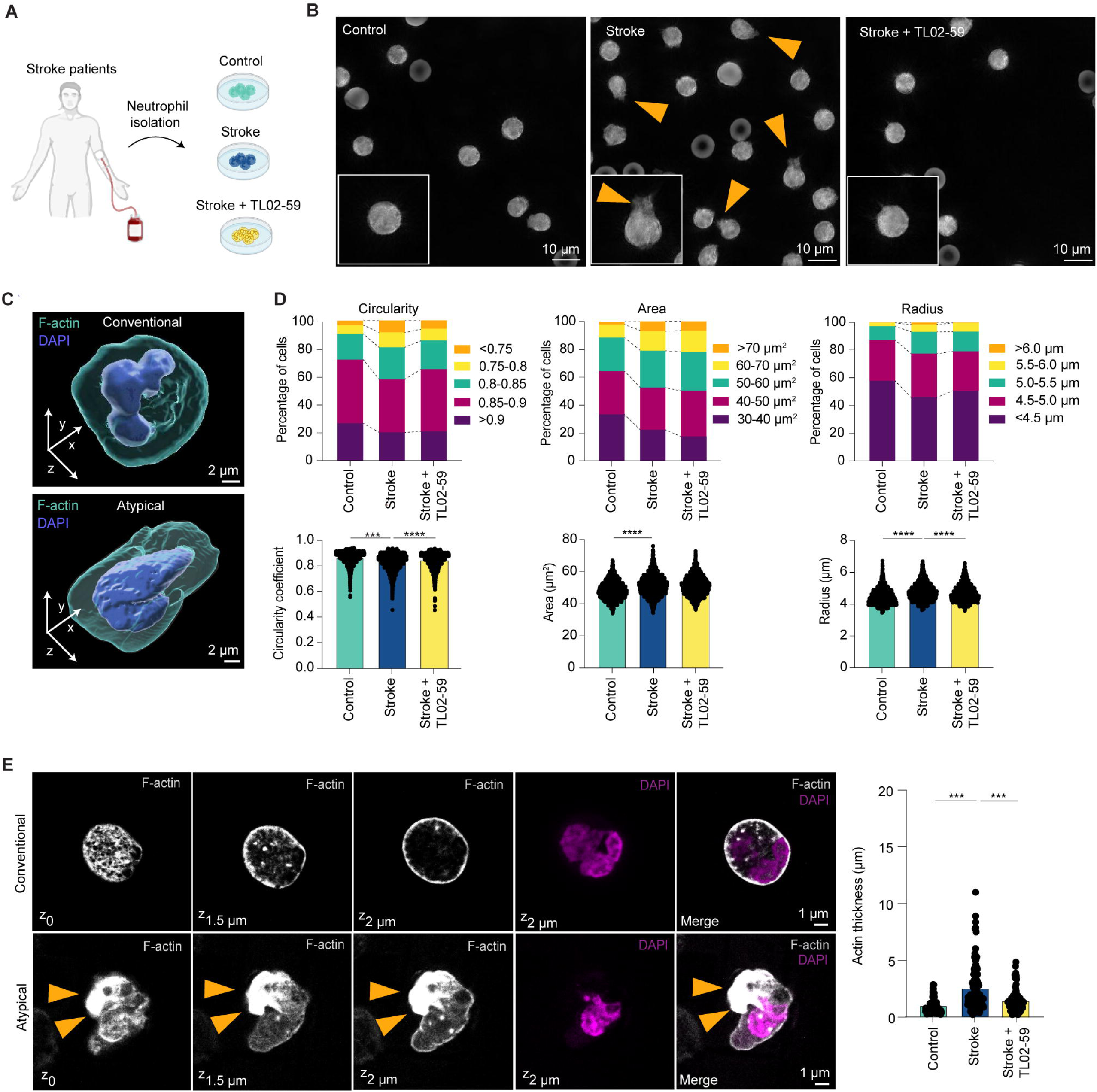
Stroke patients show an increase in atypical neutrophils, which can be reversed by pharmacological inhibition of Fgr. (**A**) Schematic for neutrophil isolation from stroke patients and healthy controls. (**B**) Representative digital holo-tomographic microscopy (DHTM) images, showing neutrophils from healthy control, stroke patients and stroke patients neutrophils treated with TL02-59. (**C**) Representative 3D images of conventional and atypical human neutrophils stained with DAPI and F-actin. (**D**) Assessment of circularity, area, and radius of neutrophils from controls (n = 1908 cells from 3 controls), stroke patients (n = 5408 cells from 4 stroke patients), and stroke patient neutrophils treated with TL02-59 (n = 3976 cells from 4 stroke patients, unpaired two-tailed t-test). (**E**) Representative immunohistological images of conventional and atypical human neutrophils. Quantifications of actin thickness in neutrophils from controls, stroke patients, and stroke patient neutrophils treated with TL02-59 (n = 298 cells from 3-4 donors per group, unpaired two-tailed t-test). Data are represented as scatter bar plots with the mean (± SEM). Statistical significance is reported as: n.s. P > 0.05, *P < 0.05, **P < 0.01, ***P < 0.001, ****P < 0.0001.

We also combined DHTM with fluorescence microscopy after staining with a nuclear dye and F-actin marker. Both fixed and live cell imaging revealed morphological heterogeneity and increased frequency of neutrophils with atypical shapes and enhanced protrusions in stroke patients, consistent with a stalling-prone phenotype (**Figure 8C,E**). We also observed an increase in neutrophils with low circularity (circularity <0.8) from 7.5% to 18.2% in the blood of patients (**Figure 8D**). These experiments suggest the presence of neutrophils with a morphology consistent with the stalling phenotype in human stroke patients, that we have observed in mice, supporting the translational relevance of our findings.

## Discussion

Here, we have identified a distinct mechanical and behavioral transition of neutrophils during stroke that critically gives rise to pathological stalling in small capillaries. This stalling-prone subset of neutrophils is characterized by altered cytoskeletal architecture, adhesive protrusions, and impaired capillary transit. This architectural heterogeneity was observed in both humans and mice. Importantly, these reprogrammed neutrophils obstruct capillaries not only within the ischemic brain but also across remote organs, thereby extending microvascular dysfunction beyond the primary lesion. In adoptive transfer experiments, stroke-derived neutrophils alone were sufficient to trigger capillary stalls and blood-brain barrier leakage in healthy recipients, demonstrating an intrinsic, transferable property of neutrophils. Importantly, this effect could be reversed by inhibition of Fgr signaling, emphasizing the therapeutic potential of our findings.

Normal organ function is critically dependent on the adequate supply of oxygen and nutrients through capillaries and maintenance of capillary flow, particularly in the brain, lungs and heart ^73–79^. Unexpectedly, we found that capillary stalls can arise from cell-intrinsic mechanisms and distinct from those occurring within the injured area, which are driven by local vascular injury. Even though recent progress has been made in understanding the mechanisms behind capillary stalls linked to pericyte dysfunction, glycocalyx degradation, and VEGF signaling ^80–82^, an important gap remains in our understanding of neutrophil-induced stalls. Identification of this cell-intrinsic mechanism demonstrates that stalls may occur independent of local vascular inflammation and may help explain the heterogeneity observed in neutrophils flow behavior. Capillary stalls are not unique to stroke and occur in neuroinflammatory conditions including Alzheimer’s disease, traumatic brain injury, and diabetes ^23,27,83,84^. Our data argue for systemic reprogramming of innate immune mechanics as a source of stalling-driving neutrophils in the circulation. Determining the relative contribution of vascular and neutrophil-dependent mechanisms to overall stalling remains an important question for future work.

Our study also unveils a widespread distribution of capillary stalls across various organs, highlighting that stroke is not a purely localized vascular event. Although stroke research has focused on neurological alterations in the brain, current research has begun to identify systemic inflammation as a critical factor affecting the short- and long-term prognosis of stroke patients ^12,85,86^. Interestingly, many of the pre-existing or acquired health conditions that arise after a stroke share common inflammatory mechanisms that can potentially exacerbate the development of other medical complications and worsen long-term outcome. Recent work has emphasized the critical role of the immune system and “whole-body macroenvironment” in shaping disease progression ^87,88^. In line with this notion, the brain continuously interacts with visceral organs through neural (via autonomic nervous system), endocrine, and immune pathways. We propose that stroke reprograms neutrophils into a pathological stalling phenotype that disseminates disease across multiple organs. Thus, post-stroke comorbidities may not be solely driven by autonomic imbalance or generalized immunosuppression but may also involve mechanical reprogramming of neutrophil transport as a possible unifying driver of multiorgan vulnerability.

On the cellular level, we connect cellular architecture to microvascular transit dynamics. The stalling neutrophil subpopulation adopted a protrusion morphology that favors the stalling phenotype. Previous studies reported that cortical actin flow drives motility and protrusion formation ^89–91^. We recapitulated this observation in neutrophils and showed that stalling cells exhibited thickened cortical actin and increased protrusions. Our data suggest that protrusions in stalling neutrophils are actin driven. Stroke-derived neutrophils exhibited enlarged areas, reduced circularity, and polarized F-actin accumulation at the cell cortex, consistent with increased protrusive stiffness that limits passage through narrow capillaries. *In vivo* imaging confirmed that stalling neutrophils lacked dynamic actin flow but instead displayed immobilized F-actin fronts coupled to adhesive microvilli, a configuration that promotes vascular arrest. These cellular alterations mirror previous observations linking integrin engagement to cytoskeletal stiffening, but here they appear to be amplified into a stable stalling state. Similarly, previous studies observed an increase in polymerized actin fibers when cells were adherent and that the actin cytoskeleton is a major driver of adhesive migration ^92,93^. Furthermore, our data complement previous studies showing that neutrophil microvilli, which are directly connected to the actin cytoskeleton, increase cellular adhesion ^94,95,96,97^. At the transcriptional level, we find enrichment of Fgr and PI3K signaling pathways, implicating Src-family kinases as upstream regulators of this mechanical program. Previous studies showed that Src kinases (Fgr, Hck and Lyn) are essential for activation of neutrophil respiratory bursts, migration, proliferation and degranulation ^98^, and that genetic deletion of these kinases impairs post-adhesion strengthening capacity, transmigration and superoxide production ^99,98,100^. In the future, it will be important to identify the location and the time of neutrophil transition into stalling states. It would be exciting to perform further studies to understand the origin of the stalling neutrophils and address many open questions in the field. For example, what are the signals or molecular cues that trigger the stalling behavior? Does passage of neutrophils through affected cerebrovascular capillaries change their state? What is the physiological (non-pathogenic) or protective function in evolution that is taken over by stroke?

We identified the Src-family kinase Fgr as a key driver of this phenotype. The increased Fgr signaling in neutrophils correlated with increased cell protrusion formation. By contrast, the conventional neutrophil subpopulation showed reduced Fgr signaling associated with small and round shapes. Fgr has been implicated in integrin signaling and cytoskeletal regulation, but its role in neutrophil capillary transit was unknown. Here, genetic and pharmacological inhibition of Fgr normalized neutrophil deformability, reduced capillary stalls, and improved tissue perfusion. Importantly, previous studies using the Fgr inhibitor TL02-59 showed that interfering with this pathway does not compromise antimicrobial defense, highlighting its therapeutic applicability ^70^. These findings elevate Fgr from a canonical signaling molecule to a potential therapeutic target for systemic complications of stroke. Our human data reinforce translational relevance. Neutrophils from stroke patients displayed the same adhesive morphology and impaired deformability as in murine models, and pharmacological inhibition of Fgr successfully reversed this phenotype ex vivo. Thus, we propose that the stalling phenotype is conserved across species and is potentially clinically actionable.

Together, our results introduce neutrophil mechanics as a mechanistic link between brain injury and systemic organ dysfunction. This conceptual shift broadens the scope of stroke therapy from local neuroprotection to systemic vascular protection, positioning neutrophil-directed interventions as a promising strategy to improve both cerebral and multiorgan outcomes.

## Limitations of the study

There are several limitations to our study. While our study focused on capillary stalls, we did not examine the functional effect of stalling on organ function. Although microcirculation failure and perfusion deficit are among the most prominent factors leading to organ injury ^101–103^, the potential impact of these capillary stalls warrants further investigation, particularly with respect to tissue oxygenation and organ damage.

Neutrophil heterogeneity in circulation and tissues is now well-recognized, with phenotype and function tightly shaped by the local tissue environment ^104,105^. Although we highlighted morphological and phenotypical changes in neutrophils as a key driver for capillary stalling, additional mechanisms, such as the release of neutrophil extracellular traps (NETs), cannot be excluded. Further studies are needed to elucidate whether NETs, which are known to promote thrombosis^106^, provide insights into peripheral pathways for generation of atypical neutrophils beyond the brain. Another unresolved question is the exact molecular mechanisms by which stroke induces expansion of a neutrophil subpopulation. Finally, we did not test the generalizability of the Fgr-mediated stalling pathway to other conditions, such as acute organ injury or infection models. Addressing these questions will be important for a comprehensive understanding of neutrophil-induced injury.

## Acknowledgements and Funding

We would like to thank Michael Sixt for the LifeAct^eGFP^ bone marrow. We would also like to thank Gabriele Wögenstein for the retina preparation. The authors acknowledge grant support from the Swiss Heart Foundation and Vontobel foundation. J.H. acknowledges funding from ANR projects “Criticality” (ANR-23-CE30-0006) and “BreakingTheWall” (ANR-22-CE35-0008).

## Author Contributions

M.E.A. and J.D. designed and performed the experiments and conducted statistical analysis. J.H., C.G., A.C.F., C.S., T.B., L.O., L.H, and B.P. performed experiments and data analysis. H.P. and L.G. performed data analysis and visualization. I.M., M.G., S.T., M.C.A., N.N., C.S., and A.H., helped with conceptualization and organization of the experiments. B.W., S.W. and M.E.A. reviewed and edited the manuscript, supervised the project and secured the needed funding. All authors were involved in critically reviewing the manuscript.

## Methods

### Key resources

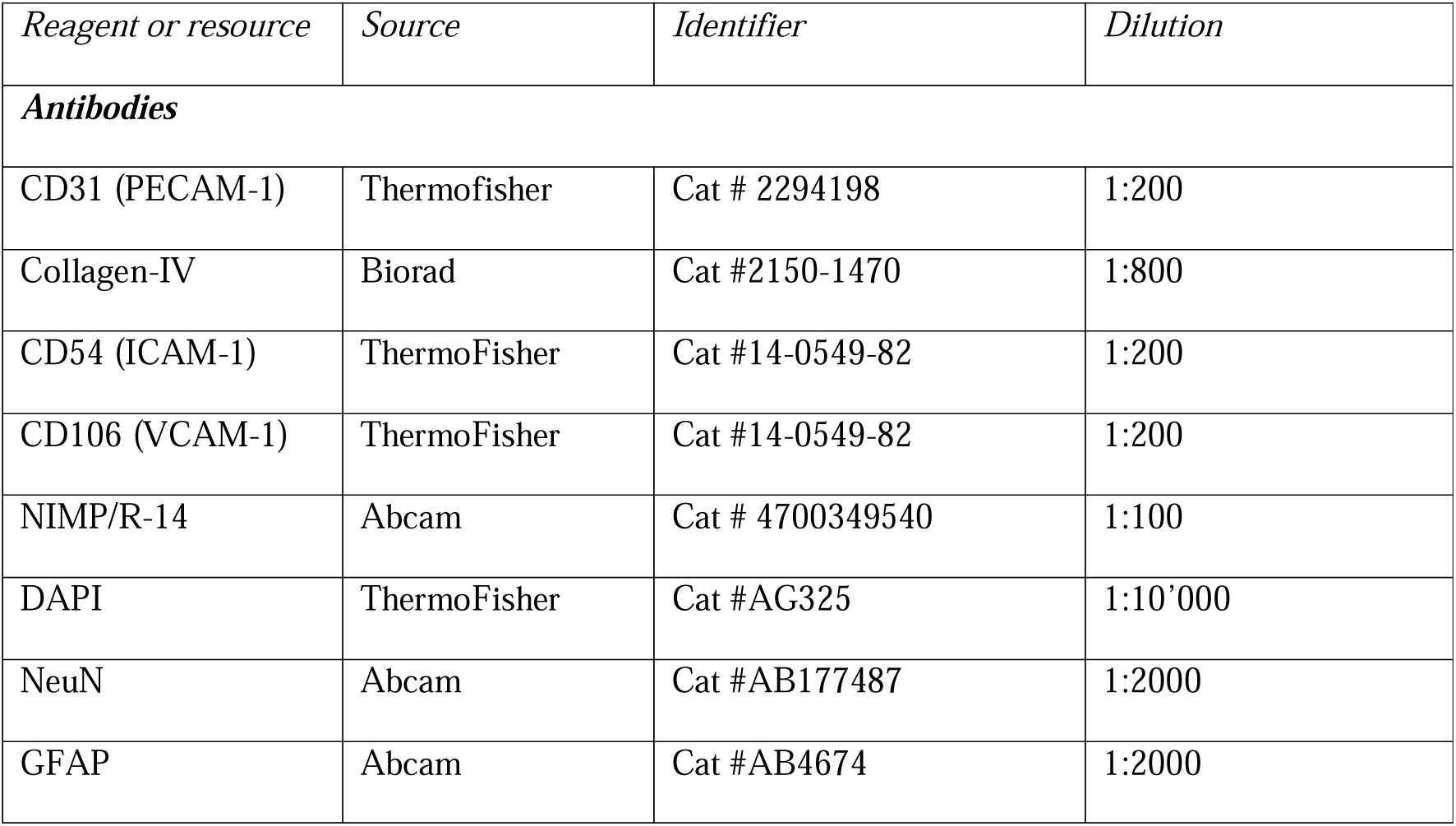
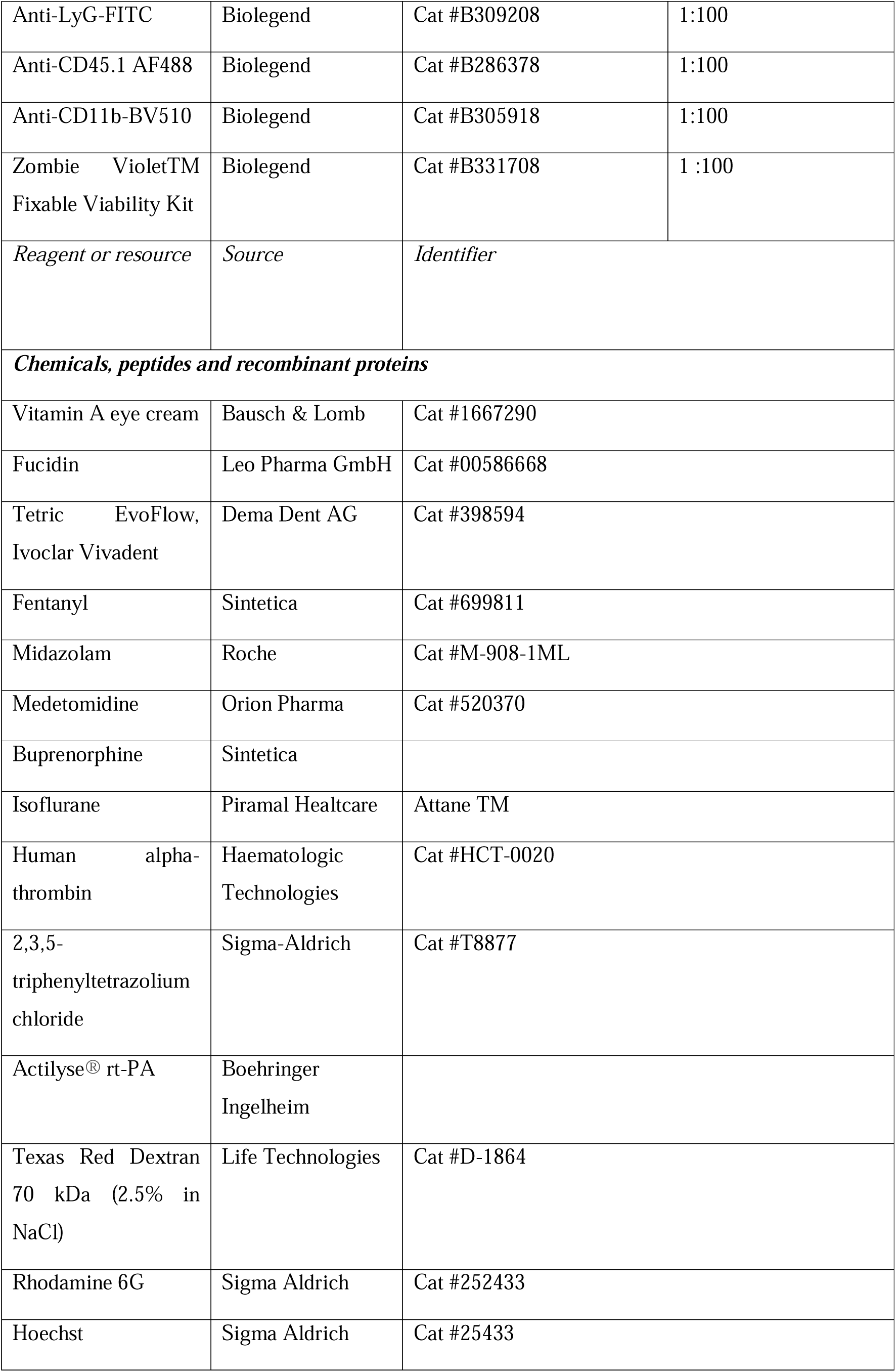
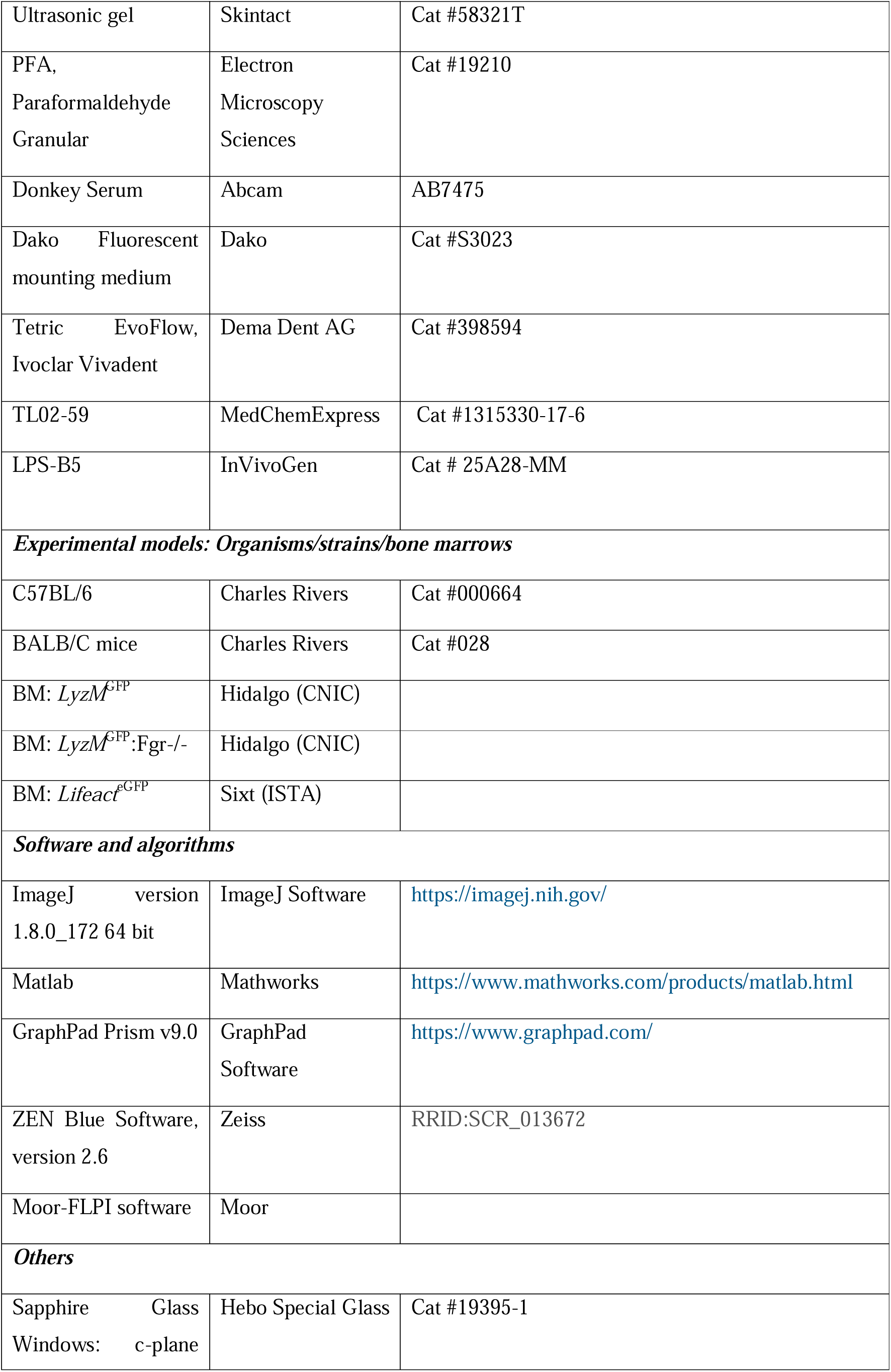
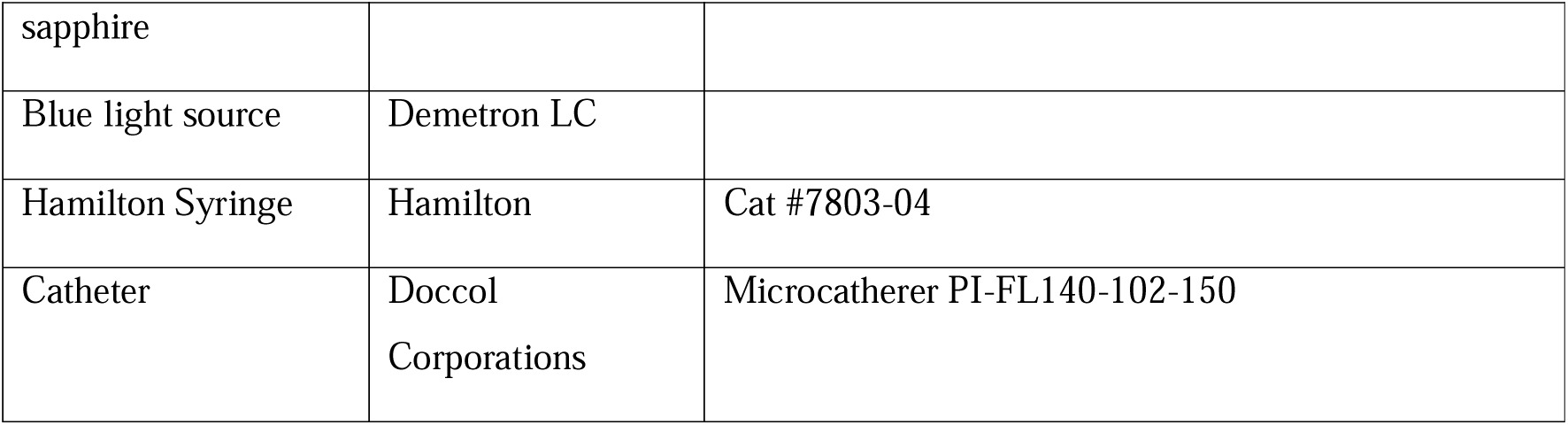

| *Reagent or resource* | *Source* | *Identifier* | *Dilution* |
| --- | --- | --- | --- |
| **Antibodies** |  |  |  |
| CD31 (PECAM-1) | ThermoFisher | Cat # 2294198 | 1:200 |
| Collagen-IV | Biorad | Cat #2150-1470 | 1:800 |
| CD54 (ICAM-1) | ThermoFisher | Cat #14-0549-82 | 1:200 |
| CD106 (VCAM-1) | ThermoFisher | Cat #14-0549-82 | 1:200 |
| NIMP/R-14 | Abcam | Cat # 4700349540 | 1:100 |
| DAPI | ThermoFisher | Cat #AG325 | 1:10'000 |
| NeuN | Abcam | Cat #AB177487 | 1:2000 |
| GFAP | Abcam | Cat #AB4674 | 1:2000 |
| Anti-LyG-FITC | Biolegend | Cat #B309208 | 1:100 |
| Anti-CD45.1 AF488 | Biolegend | Cat #B286378 | 1:100 |
| Anti-CD11b-BV510 | Biolegend | Cat #B305918 | 1:100 |
| Zombie Violet™<br>Fixable Viability Kit | Biolegend | Cat #B331708 | 1 :100 |
| *Reagent or resource* | *Source* | *Identifier* |  |
| ***Chemicals, peptides and recombinant proteins*** |  |  |  |
| Vitamin A eye cream | Bausch & Lomb | Cat #1667290 |  |
| Fucidin | Leo Pharma GmbH | Cat #00586668 |  |
| Tetric EvoFlow,<br>Ivoclar Vivadent | Dema Dent AG | Cat #398594 |  |
| Fentanyl | Sintetica | Cat #699811 |  |
| Midazolam | Roche | Cat #M-908-1ML |  |
| Medetomidine | Orion Pharma | Cat #520370 |  |
| Buprenorphine | Sintetica |  |  |
| Isoflurane | Piramal Healthcare | Attane TM |  |
| Human alpha-thrombin | Haematologic<br>Technologies | Cat #HCT-0020 |  |
| 2,3,5-triphenyltetrazolium<br>chloride | Sigma-Aldrich | Cat #T8877 |  |
| Actilyse® rt-PA | Boehringer<br>Ingelheim |  |  |
| Texas Red Dextran<br>70 kDa (2.5% in<br>NaCl) | Life Technologies | Cat #D-1864 |  |
| Rhodamine 6G | Sigma Aldrich | Cat #252433 |  |
| Hoechst | Sigma Aldrich | Cat #25433 |  |

|  |  |  |
| --- | --- | --- |
| Ultrasonic gel | Skintact | Cat #58321T |
| PFA,<br>Paraformaldehyde<br>Granular | Electron<br>Microscopy<br>Sciences | Cat #19210 |
| Donkey Serum | Abcam | AB7475 |
| Dako Fluorescent<br>mounting medium | Dako | Cat #S3023 |
| Tetric EvoFlow,<br>Ivoclar Vivadent | Dema Dent AG | Cat #398594 |
| TL02-59 | MedChemExpress | Cat #1315330-17-6 |
| LPS-B5 | InVivoGen | Cat # 25A28-MM |
| ***Experimental models: Organisms/strains/bone marrows*** |  |  |
| C57BL/6 | Charles Rivers | Cat #000664 |
| BALB/C mice | Charles Rivers | Cat #028 |
| BM: $LyzM^{GFP}$ | Hidalgo (CNIC) | |
| BM: $LyzM^{GFP}:Fgr^{-/-}$ | Hidalgo (CNIC) | |
| BM: $Lifeact^{eGFP}$ | Sixt (ISTA) | |
| ***Software and algorithms*** |  |  |
| ImageJ version<br>1.8.0_172 64 bit | ImageJ Software | https://imagej.nih.gov/ |
| Matlab | Mathworks | https://www.mathworks.com/products/matlab.html |
| GraphPad Prism v9.0 | GraphPad<br>Software | https://www.graphpad.com/ |
| ZEN Blue Software,<br>version 2.6 | Zeiss | RRID:SCR_013672 |
| Moor-FLPI software | Moor |  |
| ***Others*** |  |  |
| Sapphire Glass<br>Windows: c-plane | Hebo Special Glass | Cat #19395-1 |
| sapphire |  |  |
| Blue light source | Demetron LC |  |
| Hamilton Syringe | Hamilton | Cat #7803-04 |
| Catheter | Docol<br>Corporations | Microcatherer PI-FL140-102-150 |

### Animals

C57BL/6 and BALB/c mice (females and males, 8-12 weeks, Charles River’s) were used for this study. Animals were housed in individually ventilated, temperature-controlled cages under a 12-hour dark/light cycle. Pelleted food (3437PXL15, CARGILL) and water were provided ad-libitum. All experiments were performed in accordance with the Swiss Federal Act on Animal Protection and the ethics committee for animal research of Paris and the French ministry of higher education, research, and innovation (APAFIS #46944). Experiments were conformed to the Cantonal Veterinary Office Zurich (ZH165/2019) and (ZH030/2023) and the European Community Council Directive 2010/63/UE of September 22.2010 on the protection of animals used for scientific purposes.

### Anesthesia and preoperative monitoring

For head-post and cranial window implantation, animals were anesthetized with a subcutaneous (s.c.) injection of a mixture of fentanyl (0.05 mg/kg bodyweight), midazolam (5mg/kg bodyweight) and medetomidine (0.5 mg/kg bodyweight). Throughout the surgery, oxygen was provided to the mouse at a rate of 500 ml/min. For stroke surgery and two-photon imaging and 30 minutes before anesthesia induction, mice were injected with a s.c. dose of buprenorphine (0.1 mg/kg bodyweight). Then, anesthesia was induced with 4% isoflurane and maintained at 1.5% with continuous supply of oxygen and air (30%/70%). During all surgery and imaging sessions, the mouse body temperature was kept constant at 37° C using a thermic blanket heating system (Harvard Apparatus). The eyes were kept moist with a vitamin A eye cream.

### Head-post and cranial window implantation

Surgical interventions were performed in female and male C57BL7/6 and BALB/C at age 8-12 weeks. Animals were head-fixed in a stereotaxic frame (Model 900; David Kopf Instruments). Mice were anesthetized, the heads were shaved and the hair removed with shaving cream. The head was cleaned with surgical antiseptic solution and local anesthetics were applied subcutaneously 3 minutes before surgery (lidocaine and bupivacaine). The skin of the scalp was removed, the skull cleaned, and bonding agent was applied. The headpost was formed by multiple layers of light-curing dental cement and polymerization was done with a handheld blue light source (600 mW/cm2; Demtron LC). A custom-made aluminum head-post was attached to the head of the mouse. On the left hemisphere of the skull above the somatosensory cortex, craniotomy was performed using a dental drill (OSSEODOC; Bien-Air). The skull and dura were removed and a 3x3 mm squared sapphire glass was placed on the brain and the open areas closed with dental cement. The skin lesion was treated with an antibiotic cream (Fucidin, LEO Pharma GmbH). After surgery, animals were kept warm and received analgesics (buprenorphine 0.1mg/kg bodyweight). Mice were imaged and used for experiments after two weeks of recovery post-surgery.

### Thrombin stroke model

We used the previously described model of stroke^107,108^ for our study. In brief, mice were head-fixed on a stereotaxic frame, the fur on the neck and head was shaved and shaving cream was applied. The skin was incised with one cut in between the eyes and ears of the mice. The left temporal muscle was retracted, and a small craniotomy was drilled above the middle cerebral artery (MCA) on the M2 segment. Then, the dura was removed, and glass pipette cut by hand was introduced into the MCA. The glass pipette was filled before the surgery with human alpha-thrombin (1.5UI; HCT-0020, Haematologic Technologies Inc., USA). The alpha-thrombin was slowly injected to form a clot in situ into the lumen of the MCA. Once the clot was stable, the pipette was removed, and ischemia assessed with laser speckle imaging or laser doppler. Only a drop of at least 50% from baseline levels in the MCA territory was considered a successful stroke.

### Lipopolysaccharide (LPS) inflammation model

We injected lipopolysaccharide (1mg/kg, Invivogen) or saline as control intraperitoneally into mice and performed imaging or immunohistochemistry 12 hours post-injection.

### Stroke treatment with t-PA and Fgr administration

Thrombolysis with human rt-PA (10 mg/kg Actylyse, Boehringer Ingelheim) was performed 30 minutes after stroke induction and given through i.v. injection with a syringe pump at constant infusion speed (10% of rt-PA was given as a bolus for 30 seconds and the 90 % was infused for 30 minutes) at a rate of 6 µl/min. To inhibit Fgr, we dissolved TL02-59 (Sigma-Aldrich) in saline and injected 5 mg/kg per mouse as a bolus before rt-PA infusion. Control mice received saline. For ex vivo application of Fgr, we added 10 nM of the inhibitor (Medchem Express, Mammoth Junction, NJ, USA, Cat no: HY-112852) and incubated for 90 minutes before imaging.

### Laser speckle contrast imaging (LSCI)

Cortical cerebral blood flow (CBF) was monitored using a commercial laser speckle contrast imaging device (FLPI, Moor Instruments, UK). Mice were anaesthetized and fixed on the stereotactic frame for surgery. CBF was recorded prior to cerebral ischemia (baseline) after stroke induction as well as after rt-PA treatment after 30 minutes. CBF was analyzed from different regions of interests (ROI) as the core and penumbra of the stroke. The LSC images were generated with arbitrary units in a 16-color palette by the MoorFLPI software.

### Two-photon microscopy

The two-photon microscope consisted of a custom-built two-photon laser scanning microscope. This microscopy was equipped with a tunable pulsed laser (Chameleon Discovery TPC Coherent Inc.) and a 25x water immersion objective (W-plan Apochromat 25x/0.1). The emission of fluorophores was detected with GaAsP photomultiplier modules (Hamamatsu Photonics) and the band bass filters had a range of 475/64, 520/50 and 606/70 nm. The dichroic mirrors separated the light at 506, 560 and 652 nm (BrightLine; Semrock). The microscopes were manipulated through a customized version of ScanImage (r3.8.1; Janelia Research Campus)^109^. Echography gel was applied on the cranial window of mice for immersion of the objective. To visualize the vasculature and blood cells, Texas Red Dextran (2.5% in NaCl, 70 kDa, 50 µl), Rhodamine 6G (50µl of 1mg/ml solution in saline) and Hoechst 33342 (50 µl of 4.8 mg/ml in saline) were injected intravenously with a tail vein catheter. All dyes were injected before imaging and excited at 900 nm. The regions of interest determining the core and penumbra were taken accordingly to the LSI; the core corresponding to <30 % and penumbra corresponding to 30-50% of baseline tissue perfusion. Z-stacks were acquired with 512x512 pixels at 0.74 Hz and with 1 µm step sizes in the z-place. Z-stacks covered roughly a volume of 240 x 240 x 300 µm^3^.

### Neutrophil isolation

Whole blood was drawn via cardiac puncture and collected into EDTA-coated tubes. Neutrophils were isolated from whole blood by density-gradient centrifugation at 700 g for 30 minutes as described before^57^. The interface was washed with ACK lysis buffer and resuspended into RPMI-1640 with 10 % FBS and 1 % Streptomycin/Penicillin.

### FACS

For fluorescence activated cell sorting of neutrophil for further procedures, we lysed whole blood (BALB/c mice, females and males, 8-12 weeks, Charles River’s) with ACK lysis buffer. Blood was washed three times with PBS and cells were stained with anti-CD45.1-AF488, anti-CD11b-BV510 and anti-Ly-6G-FITC and ZombieViolet (Biolegend) for 30 minutes at 4°C. Cells were washed and cell sorting was performed with BDFACS Aria III 5L (BD Biosciences).

### Bone marrow chimeras

6–8-week-old C57BL/6 mice received a split dose of 2 x 5.5 Gy with a 24-hour interval. 4-6 hours after the second irradiation, a total of 5 x 106 cells from either LyzM^GFP^ or LyzM^GFP^:Fgr-/-, Lifeact^GFP^ bone marrow were injected intravenously. For two weeks after irradiation, mice were supplied with antibiotics in the drinking water. Chimerism was monitored in the blood by flow cytometry four weeks after reconstitution.

### Microfluidic chip

We fabricated microfluidic channels to mimic the brain capillaries in their dimensions and mechanical attributes. Initially, we employed layout Editor software to delineate the desired capillary network. We chose 6 µm as vessel diameter for our experiments. Following this, laser writing was employed to generate a photomask featuring multiple replicas of the design. Through UV photolithography, we transferred these patterns onto a silicon wafer coated with photoresist. SU8-3025 negative photoresist was employed to preserve the mask pattern on the silicon wafer. This wafer was treated with a Silane coating (Sigma-Aldrich, 440159-100□ml), and it then served as a mold for soft lithography. We utilized Polydimethylsiloxane (PDMS) SYLGARD™ 184 Silicone Elastomer Kit (Sigma-Aldrich) for this procedure. We mixed the Sylgard elastomer base with the Sylgard curing agent in a 10:1 ratio, and we poured the mixture onto the fabricated silicon wafer. After degassing, the mixture was baked in the oven at 85°C for 90 minutes. Once polymerized, we detached the PDMS from the wafer, cut the various designs into individual square slabs, and punched two holes for inlet and outlet. Plasma treatment was applied to each slab and a glass slide before bonding them together. The assembled device was baked in the oven at 85°C for another 90 minutes, making it ready for experimental use. For the experiments, a syringe pump was connected to the microfluidic device using silicone tubing with an inner diameter of 0.5 mm. A 1 mL syringe pre-attached to the silicone tubing was used to manually aspirate the cell suspension, after which it was incorporated into the pump system. A constant flow of 0.05 µl/s was applied during the experiments.

### Micropipette force probe

Cell mechanical properties were quantified by measuring the cell adhesion force and stiffness (Young’s modulus). Isolated neutrophils were place in a petri dish filled with RPMI 1640, 10% FCS and 1% penicillin-streptomycin at room temperature. The cells were picked up by using a rigid micropipette and pressed against a flexible micropipette with a microbead at its tip (used as a microindenter) with the help of motorized micromanipulators. Cells were then pulled away from the microindenter’s tip until the cell-tip adhesion was broken. The position of the tip of the microindenter was measured by image acquisitions with an inverted microscope, equipped with differential interference contrast (DIC), 100x objective (Plan Fluor, 1.3 numerical aperture, DIC, oil). The deflection of the bead was determined and analyzed with a customized MATLAB code^66^.

### Sequencing and data processing

After cell specific isolation with FACS, we used the commercially available RNA isolation kit. We used the RNeasy Mini Kit (QIAGEN^TM^) for RNA extraction according to manufacturer’s direction. RNA was quantified spectrophotometrically (Nanodrop, Thermo Scientific, Wilmington, DE). We used Illumina Smartseq II for RNA sequencing. Library construction and RNA sequencing was performed at the Functional Genomics Center Zurich (Zurich, Switzerland).

### Adoptive cell transfer

For adoptive cell transfer, we isolated neutrophils from healthy controls and stroke mice (24h post-stroke). We injected 300’000 neutrophils per mouse into a healthy receiver with cranial window. We injected the cells suspended in PBS with a catheter (Doccol Corporation, Sharon Massachussettes, USA) sutured into the intracarotid artery.

### Quantification of lesion volumes

The stroke lesion of mice was quantified 7 days post-stroke. Mice were euthanized through an i.p. injection and overdose of pentobarbital (200mg/kg). Lesions were assessed with 2,3,5-triphenyltetrazolium chloride (TTC, cat. # T8877, Sigma-Aldrich) or immunohistochemistry. For TTC quantifications, brains were cut directly after extraction into 1 mm thick coronal slices and put for 10 minutes into 2% TTC at 37° C. Sections were imaged and the white infarct tissue was delineated from the red tissue with an image analysis software (ImageJ, version 1.41). To calculate the total infarct volume in cubic millimeters, linear integration of the lesion areas was measured as previously described ^110^.

### Immunohistochemistry

To perform immunohistochemistry on brains and organs, mice were euthanized at day 1 or 7 post stroke. Mice received an overdose of pentobarbital (200 mg/kg) and were perfused with 50 ml PBS and 50 ml 2% PFA. The brain and organs were extracted and placed into 4 % of PFA for 4 hours at 4° C. Then, the brains and organs were placed in 30% sucrose for 3 days until frozen with dry ice. Frozen brains and organs were cut in 40 µm thin slices with a microtome (Hyrax KS 34) and stored in antifreeze solution (50 mM sodium phosphate buffer pH 7.4, 1 M glucose, 35% ethylene glycol and 3.5 mM sodium azide) at -20° C. The brain or organ sections were first blocked in 0.3% Triton X-100 and Tris-buffer (50 mM, pH 7.4) and 5% donkey serum for 1 hours at RT. Then, sections were incubated in primary antibodies diluted in blocking solution overnight at 4° C. Then, sections were incubated with secondary antibodies for 45 minutes at RT. The sections were mounted on SuperFrost Plus slides (Thermo Scientific) in Dako Fluorescence Mounting Medium (Dako, Jena, Germany). For lesion quantifications with immunohistochemistry, single sections being 1 mm apart from each other covering the whole brain were selected. Neurons were stained with anti-NeuN and nuclei with DAPI to measure the size of the lesion in mm^2^. The sum of the lesions (mm^2^) was multiplied by 1 mm to integrate the lesion in volume in cubic milimiters (mm^3^).

### Confocal microscopy

Confocal images were taken with a Zeiss LSM800 confocal laser scanning microscope. We used a 10x (Plan-Apochromat, NA 0.45) or 40x objective (Plan-Apochromat, NA 1.4, Oil DIC (UV) VIS-IR). Overview sections were taken with the 10x objective and the tile scan option from the built-in microscope software (ZEN software, Zeiss), and single stacks were taken with the 40x objective.

### Motor scoring and behavioral analysis

To assess stroke outcome, severity and recovery at days 1, 3, 5, 7 we performed several sensorimotor tests. We used small sticky tapes of approximately 5 x 5 mm for the adhesive tape removal test. The left and right forepaws of mice taped with the small sticky tapes and time to sense and remove was measured. First, animals we trained two consecutive days before stroke surgery to sense and remove the sticky tape faster than 10 seconds. After stroke surgery, on day 1, 3, 5 and 7, sticky tapes were placed on the forepaws of mice and time for sensing and removal were measured three times and averaged^111^. For assessment of the neurological score (neuroscore) we used as previously described a combination of sensorimotor functional tests^112^. The different tests consist of measuring the thorax torsion (0, 1, 2 points), forearm torsion (0, 1, 2 points), grasping reflex of the fore-and hindpaw (0, 1 point each), leg hanging test of fore-and hindpaw (0,1 point each). This summarizes to a maximal score of 8, meaning no sensorimotor deficit and deficits for lower values.

For the pole test, mice were placed on the tip of a 50 cm long wooden rod with 1 cm of diameter. The mice needed to turn and use their tail to descend the rod. The time needed to turn (T_turn_) and time to descend (T_total_) was measured. Mice were trained two consecutive days before stroke surgery. The average of 5 rounds was taken for training and performing the task post-stroke^113^.

With the rotarod test, we evaluated the mouse equilibrium and locomotion ability post-stroke as previously described^113^. The mouse was placed on a rod combined to an electric motor for rotation. The rotor was increased from 300 rpm per minute to 800 rpm within five minutes and the time each mouse could stay on the rod before falling was measured.

### Human Neutrophils Sample Preparation

The study was approved by the Cantonal Ethics Committee of Zürich (number: 2024-01353) and followed the principles of the Declaration of Helsinki. All patients provided written informed consent. Whole blood was freshly obtained from healthy controls and stroke patients at the University Hospital Zürich and collected in 10 mL K2EDTA vacutainers. Whole blood was stored at 4°C and processed within 2 hours after collection. Neutrophils were isolated from whole blood using the MACSxpress® Whole Blood Neutrophil Isolation Kit, human (Miltenyi Biotec, Germany) with the MACSxpress® Separator Kit (Miltenyi Biotec, Germany). A total of 1.8 mL of isolated neutrophils in solution was obtained from 4 mL of whole blood. Isolated neutrophils were stored at 4°C and imaged within 3 hours.

### Label-Free Digital Holo-tomographic Microscopy

Label-free holo-tomographic imaging (DHTM) was performed using a 3D Cell Explorer® microscope (Nanolive SA, Switzerland)^114^. First, isolated neutrophils in solution were further diluted 1:1 in Hanks’ Balanced Salt Solution (HBSS) (Thermo Fisher Scientific, 14025092) before imaging. For each measurement, 250 μL of the isolated neutrophil solution was transferred in a 35 mm Ibidi Bioinert μ-Dish (Ibidi GmbH, Germany) for DHTM imaging. The Petri dish was placed in the microscope sample holder, and neutrophils were allowed to sediment to the bottom of the Petri dish for 10 min before imaging. For each sample, ∼120 refractive index (RI) images were acquired containing between 3-15 single neutrophils. Each RI image corresponds to a field of view of 90 μm × 90 μm × 30 μm. The DHTM was operated under standard laboratory conditions.

For the TL02-59-treated neutrophils, 50 μL of 60 nM TL02-59 stock solution was added to 250 μL of the isolated neutrophil solution and incubated for 90 minutes at 4°C. Following, the 300 μL solution was transferred in a 35 mm Ibidi Bioinert μ-Dish (Ibidi GmbH, Germany) and imaged as described above.

### Image processing, data analysis and statistical analysis

Z-stacks and frame scans were acquired with two-photon imaging. The data were analyzed using ImageJ (NIH, version 1.41). To quantify the number of flowing, stalling, adhering and crawling neutrophils in different regions of the brain, the automated tracking software Trackmate^115^ as a Fiji plugin was used. Masks were applied with ImageJ to select for the vessels and using the Laplacian of Gaussian filter to detect the cells.

3D RI stacks obtained by DHTM were exported as TIFF files using Steve® software v1.6 (Nanolive SA, Switzerland). For the structural analysis and quantification of neutrophils, 3D RI stacks obtained by DHTM were imported into the open-source software FIJI and a maximum intensity Z-projection was applied. The 2D images were exported as TIFF files and processed using a combination of Ilastik^116^, Python-based analysis and FIJI. First, the 2D images were loaded into Ilastik, an interactive machine-learning-based image segmentation tool. A pixel classification workflow was trained to differentiate between cells and background using manually labelled training data. Following segmentation, a probability map was generated and exported as an HDF5 file. The 2D images and corresponding probability maps were then loaded into an object classification workflow to differentiate between cell types (“neutrophils”, “red blood cells”, “others”). Object prediction maps, containing only the “neutrophils” channel, were generated and exported as HDF5 files. Further downstream analysis was performed using Python in a Jupyter Notebook environment and FIJI. The object prediction mask was thresholded to create a binary mask. Each object was labelled, and the morphological properties were extracted using the analyze particles function in FIJI. The cell area, circularity and radius were extracted for each neutrophil.

Data in groups were tested for significance (p<0.05) using a student’s t-test or with the Mann-Whitney test for non-normally distributed data. Statistical analysis was performed with GraphPad Prism (version 9.0; GraphPad Software La Jolla, CA, USA). All statistical tests and the size of each group are indicated in the figure legends. Results were expressed as mean ± standard error of mean (S.E.M.).

