## Supplemental Figure for "Reprogrammed neutrophils with impaired transit mechanics drive multi-organ capillary stalling after stroke"

**Supplementary Figures:**

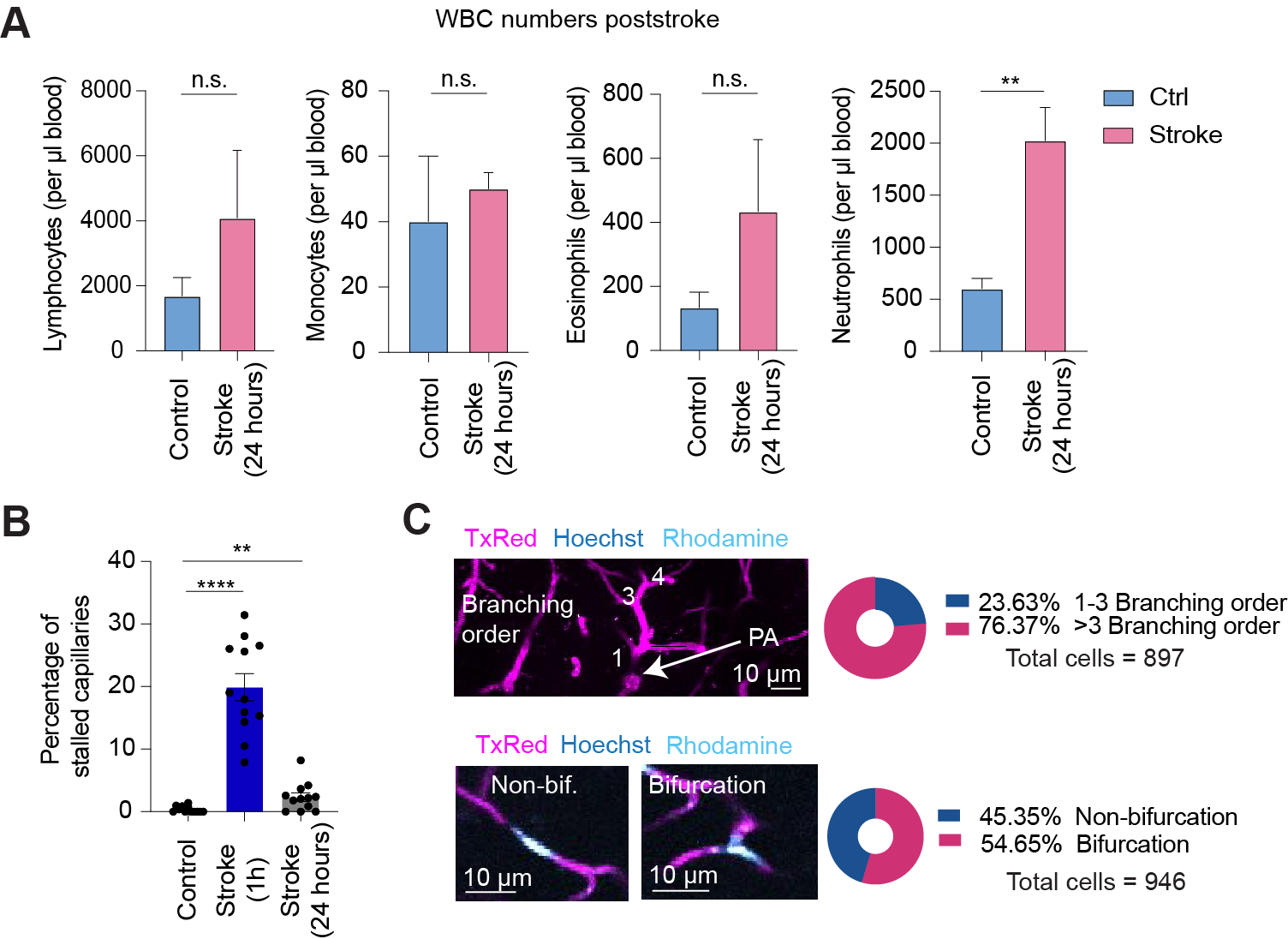

**Suppl. Figure S1: Blood circulating neutrophil numbers and capillary stalls caused by neutrophils is increased 24 hours post-stroke.**

(**A**) White blood cell (lymphocytes, monocytes, eosinophils, neutrophils) numbers in control and stroke mice blood samples, 24 hours post-stroke (n = 3 mice per group, P < 0.01, unpaired two-tailed t-test).

(**B**) Quantifications of capillary stalls by neutrophils in percentage at baseline, post-stroke and post-stroke 24 hours later (n = 12 stacks from 4 mice, P < 0.0001, unpaired two-tailed t-test).

(**C**) Neutrophil stalling types in capillaries, showing stalls in capillary birfucations and non-bifurcations and their corresponding branching orders. The different stalling types are shown as pie chart (n = 897-946 neutrophils).

Data are represented as scatter bar plots with the mean (± SEM), pie charts or bar plots with mean (± SEM). Statistical significance is reported as: n.s. P > 0.05, *P < 0.05, **P < 0.01, ***P < 0.001, ****P < 0.0001.

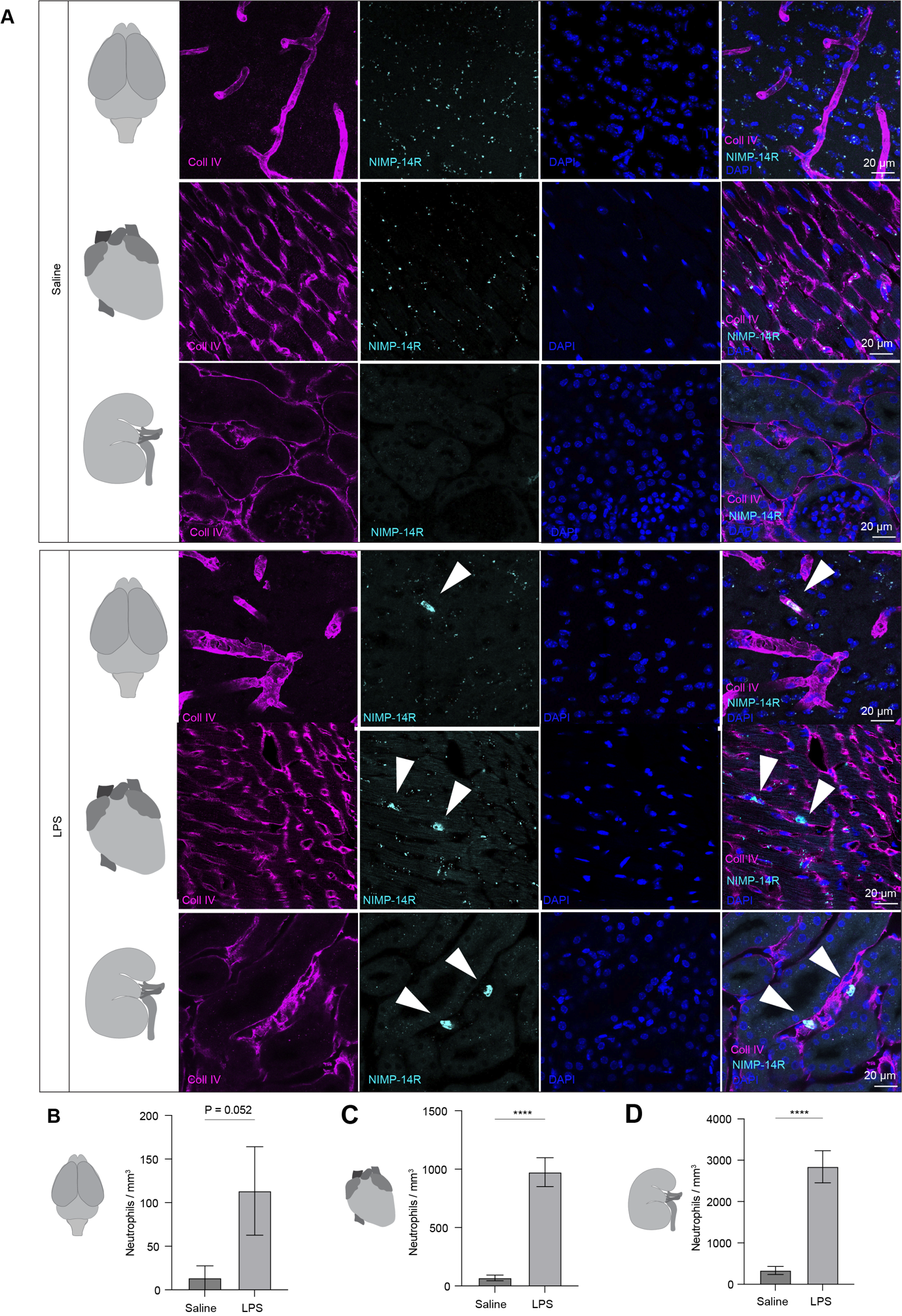

**Suppl. Figure S2: LPS injection causes an increase of vascular neutrophils in different organs.**

**(A)** Representative images of immunohistological stainings of brain, heart, and kidney after saline or LPS (1mg/kg) injection after 12 hours (n = 3 mice per group). Stained with Collagen IV, NIMP-14R and DAPI.

(**B**) Quantifications of neutrophils in the brain after saline or LPS injection (n = 12 from 3 mice per group, P > 0.05, unpaired two-tailed t-test).

(**C**) Quantifications of neutrophils in the heart after saline or LPS injection (n = 12 from 3 mice per group, P < 0.0001, unpaired two-tailed t-test).

(**D**) Quantifications of neutrophils in kidneys after saline or LPS injection (n = 12 from 3 mice per group, P < 0.0001, unpaired two-tailed t-test).

Data are represented as bar plots with the mean (± SEM). Statistical significance is reported as: n.s. P > 0.05, *P < 0.05,
**P < 0.01, ***P < 0.001, ****P < 0.0001.

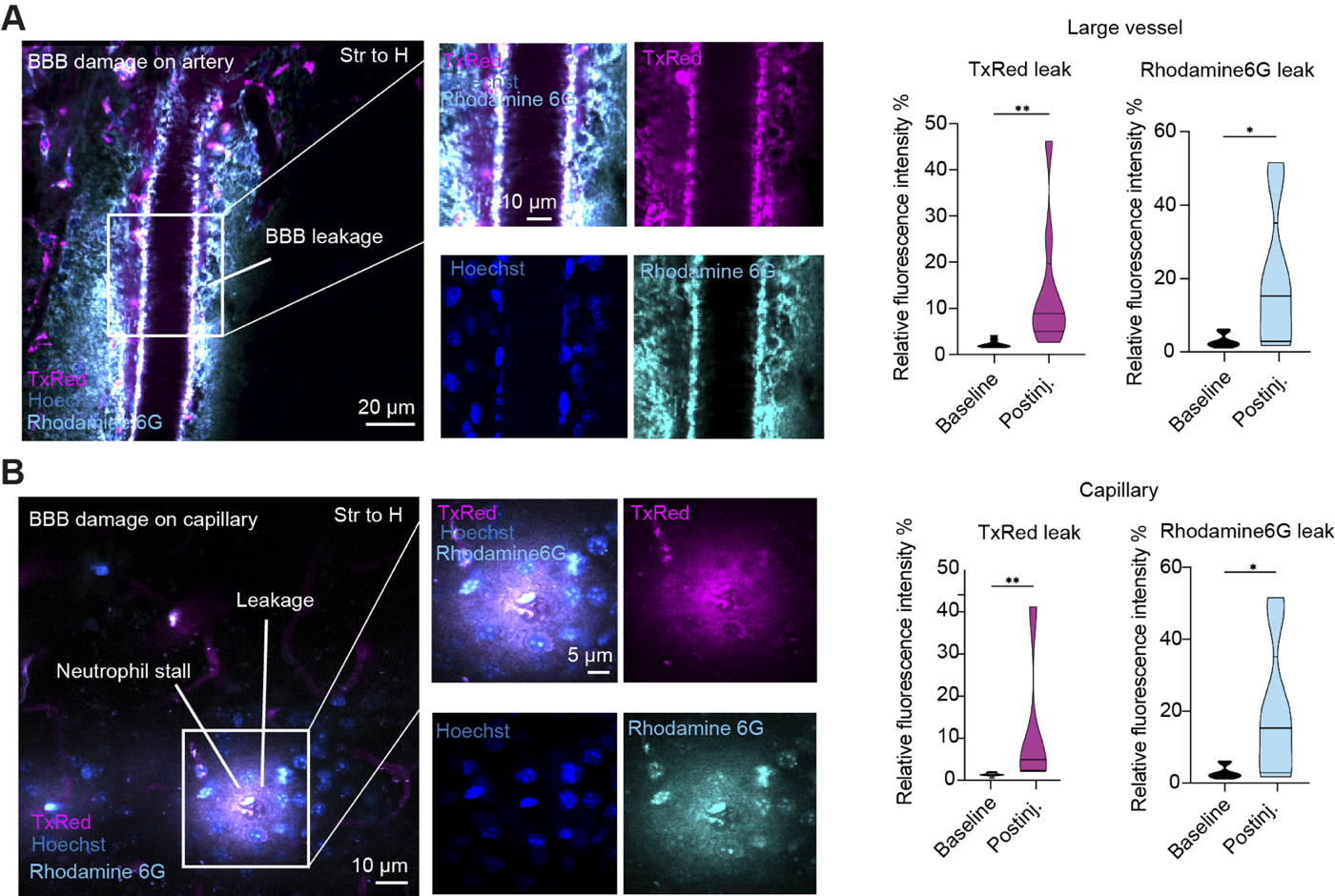

**Suppl. Figure S3: Adoptive cell transfer of stroke-derived neutrophils causes BBB leakage in healthy receiver mice.**

**(A)** Two-photon representative image of leaky large vasculature after injection of stroke-derived neutrophils, labeled with Hoechst and Rhodamine 6G. Plasma is labeled with Texas Red Dextran 70 kDa. Violin plots represent the relative fluorescence intensity of Texas Red Dextran (n = 11-12 stacks from 4 mice per group, P < 0.01, unpaired two-tailed t-test) and Rhodamine 6G (n = 11-12 stacks from 4 mice per group, P < 0.05, unpaired two-tailed t-test).

(**B**) Two-photon representative image of leaky capillaries after injection of stroke-derived neutrophils, labeled with Hoechst and Rhodamine 6G. Plasma is labeled with Texas Red Dextran 70 kDa. Violin plots represent the relative fluorescence intensity of Texas Red Dextran (n = 11-12 stacks from 4 mice per group, P < 0.01, unpaired two-tailed t-test) and Rhodamine 6G (n = 11-12 stacks from 4 mice per group, P < 0.05, unpaired two-tailed t-test).

Data are represented as violin plots showing the median and upper and lower quartiles. Statistical significance is reported as: n.s. P > 0.05, *P < 0.05, **P < 0.01, ***P < 0.001, ****P < 0.0001.

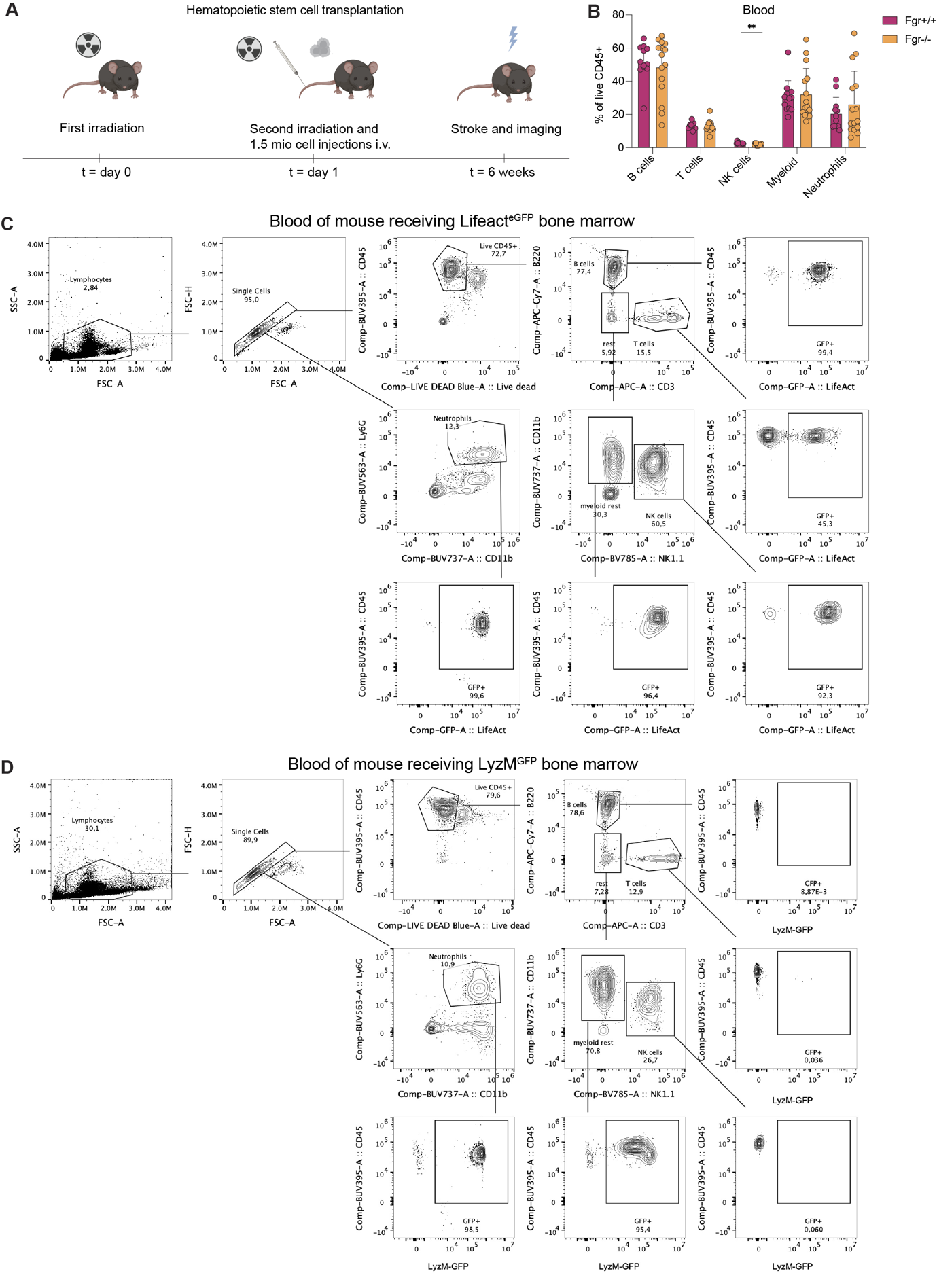

**Suppl. Figure S4: Hematopoietic stem cell transplantation of LifeAct^eGFP^, LyzM^GFP^ and LyzM^GFP^;Fgr-/- cells into receiver mice.**

**(A)** Schematic for hematopoietic stem cell transplantation.

(**B**) Recomposition of blood circulating cells (B-cells, T cells, NK cells, myeloid cells, neutrophils, n = 12-15 per group) after 6 weeks transplantation in % of live CD45 cells (NK cells P < 0.01, unpaired two-tailed t-test).

(**C**) Representative FACS gating strategy to quantify the percentage of blood cell numbers for the LifeAct^eGFP^ mice.

(**D**) Representative FACS gating strategy to quantify the percentage of blood cell numbers for the LyzM^GFP^ mice.

Data are represented as scatter bar plots showing the mean (± SEM). Statistical significance is reported as: n.s. P > 0.05, *P < 0.05, **P < 0.01, ***P < 0.001, ****P < 0.0001.

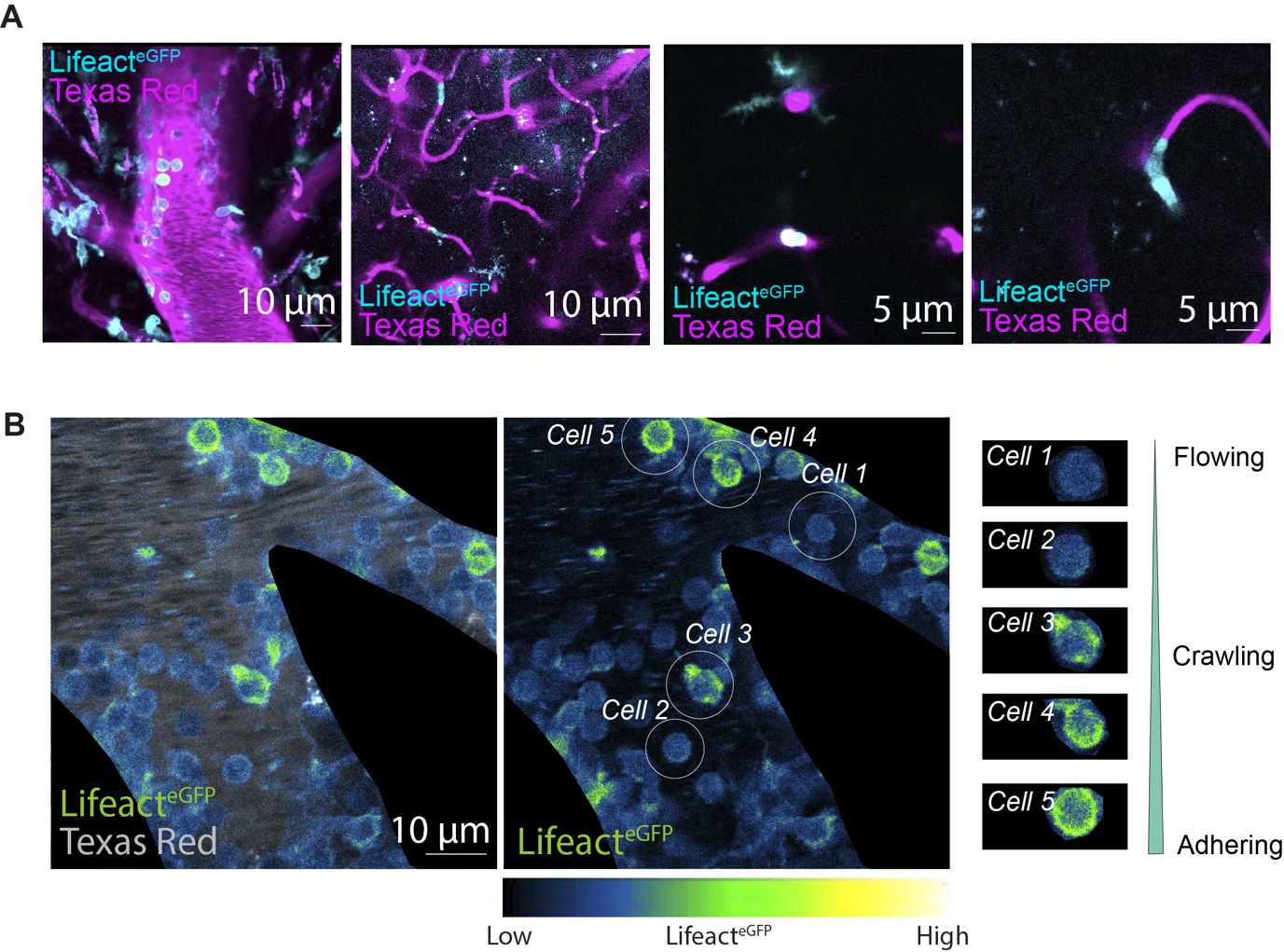

**Suppl. Figure S5: Lifeact^eGFP^ *in vivo* two-photon microscopy reveals actin dynamics in neutrophils during different cell behaviours.**

**(A)** Two-photon representative image of the brain vasculature in LifeAct^eGFP^ mice, and plasma labelled with Texas Red Dextran.

(**B**) Representative images of neutrophils from LifeAct^eGFP^ mice on large vessel while crawling, adhering and flowing.

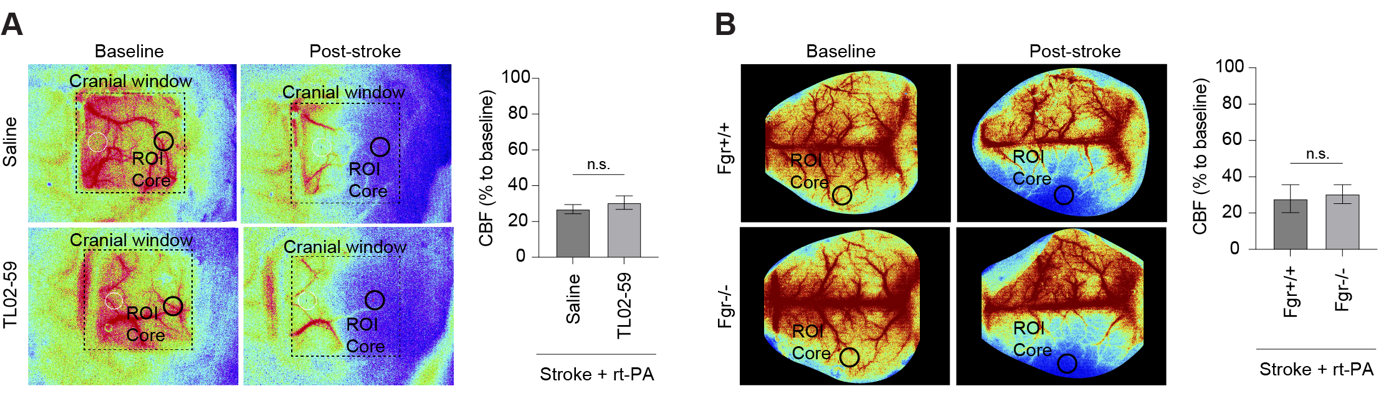

**Suppl. Figure S6: Laser speckle contrast imaging shows consistent cerebral blood flow drop post-stroke in different groups.**

**(A)** Representative images of LSCI acquisitions at baseline and poststroke with cranial windows of mice treated rt-PA and saline or rt-PA and TL02-59. Bar plots showing cerebral blood flow (CBF) drop in core region (CBF % from baseline, n = 4-5 mice per group, P > 0.05, unpaired two-tailed t-test).

(**B**) Representative images of LSCI acquisitions of Fgr+/+ and Fgr-/- mice at baseline and post-stroke. Bar plots showing cerebral blood flow (CBF) drop in core region (CBF % from baseline, n = 2-4 mice per group, P > 0.05, unpaired two-tailed t-test).

Data are represented as bar plots showing the mean (± SEM). Statistical significance is reported as: n.s. P > 0.05, *P < 0.05, **P < 0.01, ***P < 0.001, ****P < 0.0001.
